# Nonsterile immunity to cryptosporidiosis in infants is associated with mucosal IgA against the sporozoite and protection from malnutrition

**DOI:** 10.1101/2021.03.04.433864

**Authors:** Mamun Kabir, Masud Alam, Uma Nayak, Tuhinur Arju, Biplob Hossain, Rubaiya Tarannum, Amena Khatun, Jennifer A. White, Jennie Z. Ma, Rashidul Haque, William A. Petri, Carol A. Gilchrist

## Abstract

We conducted a longitudinal study of cryptosporidiosis from birth to three years of age in an urban slum of Dhaka Bangladesh. Fecal DNA was extracted from monthly surveillance samples and diarrheal stool samples collected from 392 infants from birth to three years. A pan-Cryptosporidium qPCR assay was used to identify sub-clinical and symptomatic cryptosporidiosis. Anthropometric measurements were collected quarterly to assess child nutritional status. 31% (121/392) of children experienced a single and 57% (222/392) multiple infections with *Cryptosporidium*. Repeat infections had a lower burden of parasites in the stool (Cq slope = −1.85; p<0.0001) and were more likely to be sub-clinical (Chi square test for trend; p=0.01). Repeat infections were associated with the development of growth faltering (Pearson correlation = −0.18; p=0.0004). High levels of fecal IgA antibodies against the *Cryptosporidium* Cp23 sporozoite protein at one year of life were associated with a delay in reinfection and amelioration of growth faltering through three years of life (HAZ IgA high responders −1.323 ± 0.932 versus HAZ −1.731 ± 0.984 p=0.0001). We concluded that nonsterile immunity to cryptosporidiosis in young children was associated with high levels of mucosal IgA anti-Cp23 and protection from diarrhea and growth faltering.

**Authors Summary:** *Cryptosporidium* is one of the top causes of diarrhea and growth faltering in Bangladesh infants. We discovered that a prior infection resulted in incomplete immunity that protected from diarrhea and growth faltering but not infection and was associated with mucosal IgA against a sporozoite surface protein Cp23. The most important implication of these findings is that a cryptosporidiosis vaccine may not need to achieve complete protection from infection to have a beneficial impact on child health.

## Introduction

*Cryptosporidium spp*. parasites are leading causes of diarrheal disease in infants living in low and middle income countries [1–4]. They are additionally a cause of water-borne outbreaks of diarrhea in high income countries and of chronic diarrhea in people living with HIV infection [5]. There is no vaccine and development will require an understanding of the natural history of cryptosporidiosis [3,6–12]. To this end, a community-based prospective cohort study of cryptosporidiosis was begun in 2014 [13]. The study subjects were born in an urban slum in Dhaka, Bangladesh and enrolled during the first week of life [13–15]. In humans and in animal models vaccination or prior infection resulted in partial protection against reinfection [16–19].

For example we observed that high fecal IgA against the sporozoite protein Cp23 delayed but did not prevent a repeat infection with *Cryptosporidium spp.* [20, 21]. Ajjampur et al observed a decrease in the incidence of diarrhea in reinfected children [22]. In contrast Kattula et al found that while the reinfection frequency was decreased the proportion of symptomatic disease was unchanged [9]. In human volunteer studies second infections were associated with reduced parasite burden and less severe diarrhea [23].

In addition to diarrheal disease cryptosporidiosis is associated with development of malnutrition [8,24–27]. Here we report the natural history of cryptosporidiosis from a longitudinal study of urban slum children from birth through three years of age in Dhaka, Bangladesh, demonstrating that immunity is characterized by protection from diarrhea and growth faltering.

## Results

Five hundred children were enrolled within the first week of birth, and of these 392 completed three years of observation. Stool samples were collected monthly and at the time of diarrhea. Successful sample collection and qPCR testing was completed for 96% of monthly surveillance time points and for 84% of the diarrheal cases (Fig 1; S1 Table 1). There were 1336 *Cryptosporidium* positive samples for analysis by year 3 (Fig 1). Six hundred and ninety eight events met the definition of separate *Cryptosporidium* infections in the 392 children (Table 1). Of the 698 infections experienced by the 392 infants retained in the study at 3 years of age, 167 were diarrheal and 531 sub-clinical cryptosporidiosis (Table 1). The Cq (cycle of quantification) value of the stool sample in which the parasite was first detected was used as an index of parasite burden.

**Fig 1.**
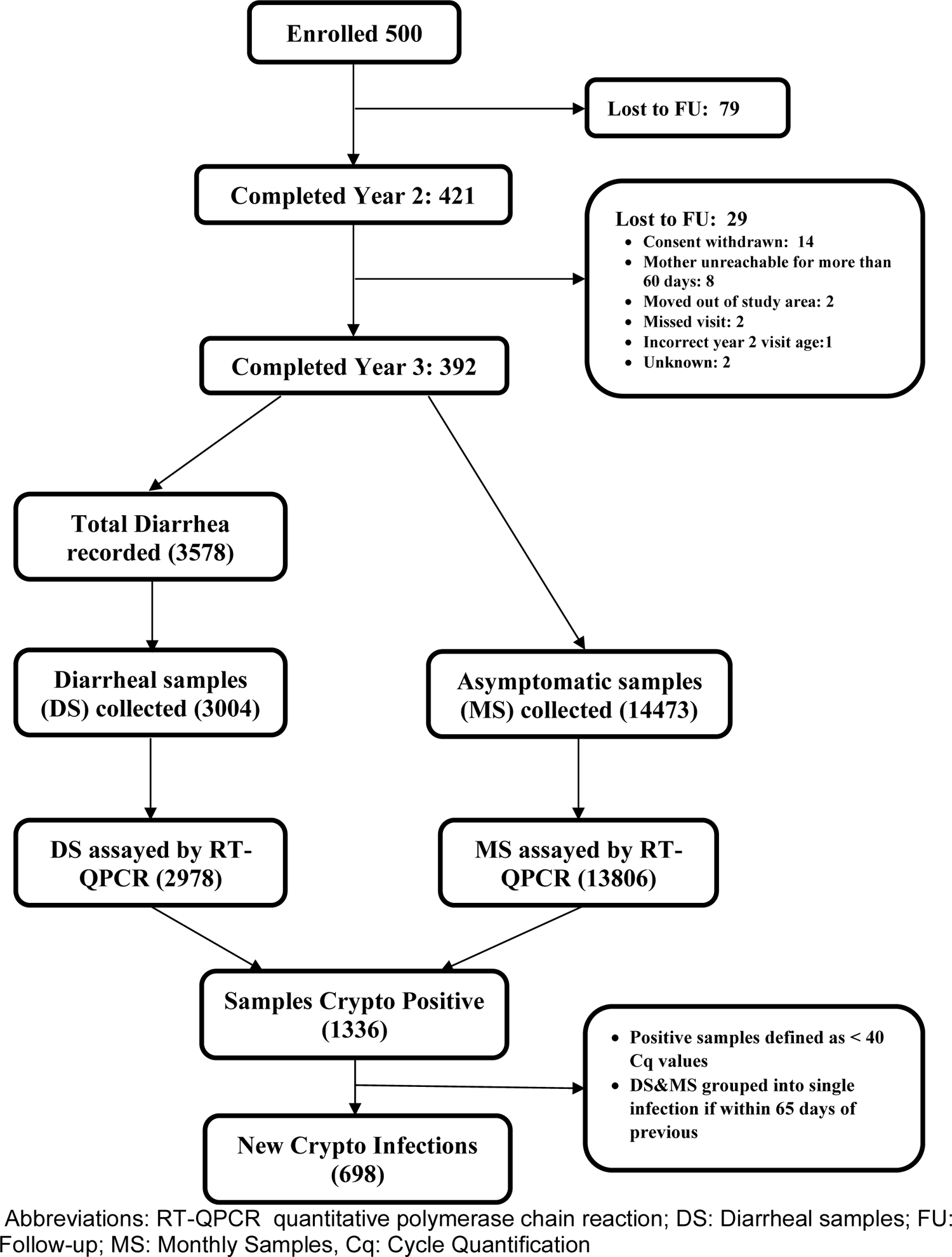
**COHORT diagram**. Study subjects, collected samples and new infection numbers

**Table 1.**
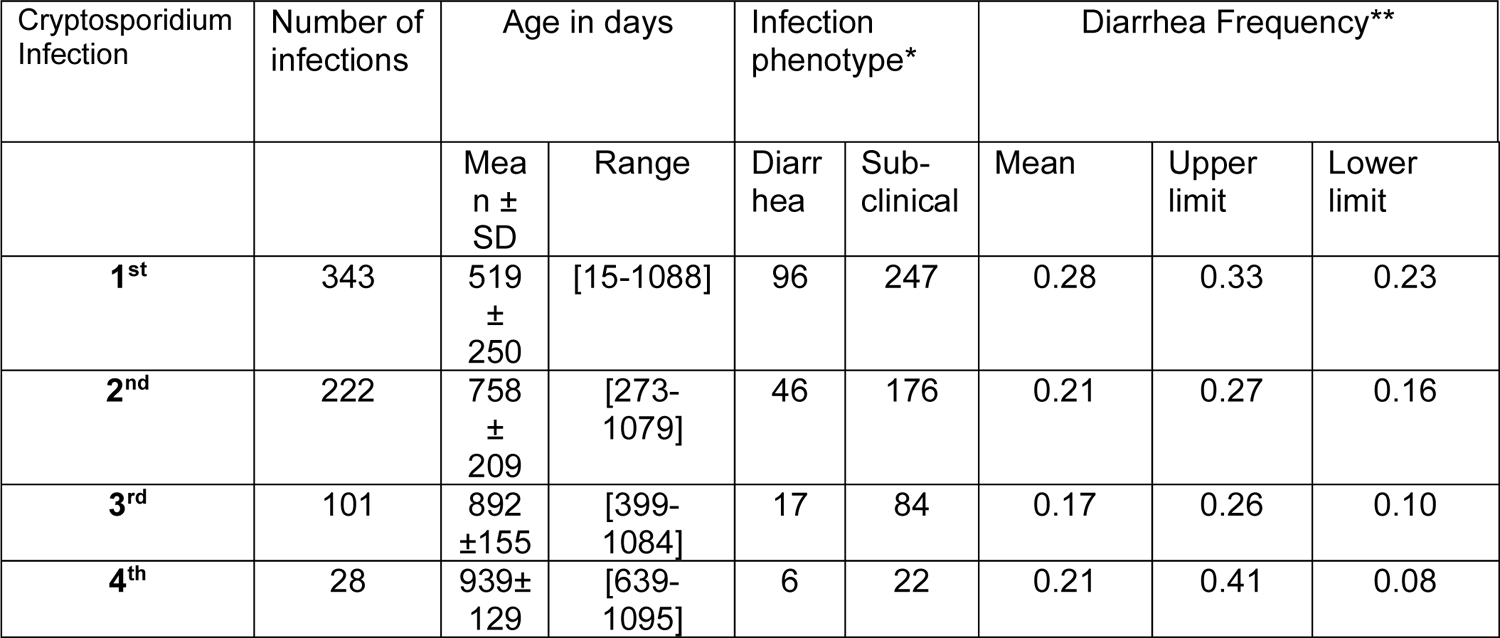
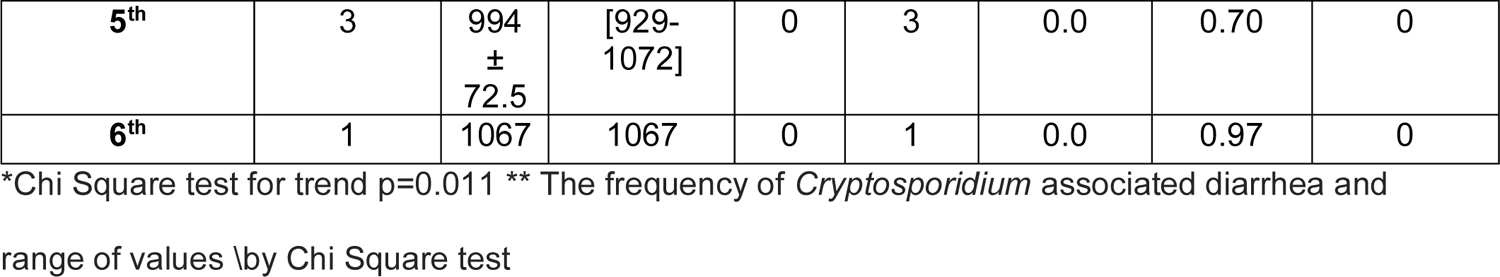
Frequency of Diarrheal Cryptosporidiosis in Repeated Infections

To investigate if the reduced severity of clinical disease in recurrent infections could be correlated with a reduction in parasite burden, the Cq values of sub-clinical and diarrheal disease were measured (Table 2). The slopes derived from the GEE models for sub-clinical (1.9 ± 0.2) and diarrheal (1.49 ± 0.31) infections were not significantly different from each other (Fig. S4). However recurrent infections had as expected a lower amount or burden of *Cryptosporidium* than did the first infection (slope −1.85 ± 0.21; p<0.0001) (Fig 2A).

**Fig 2.**
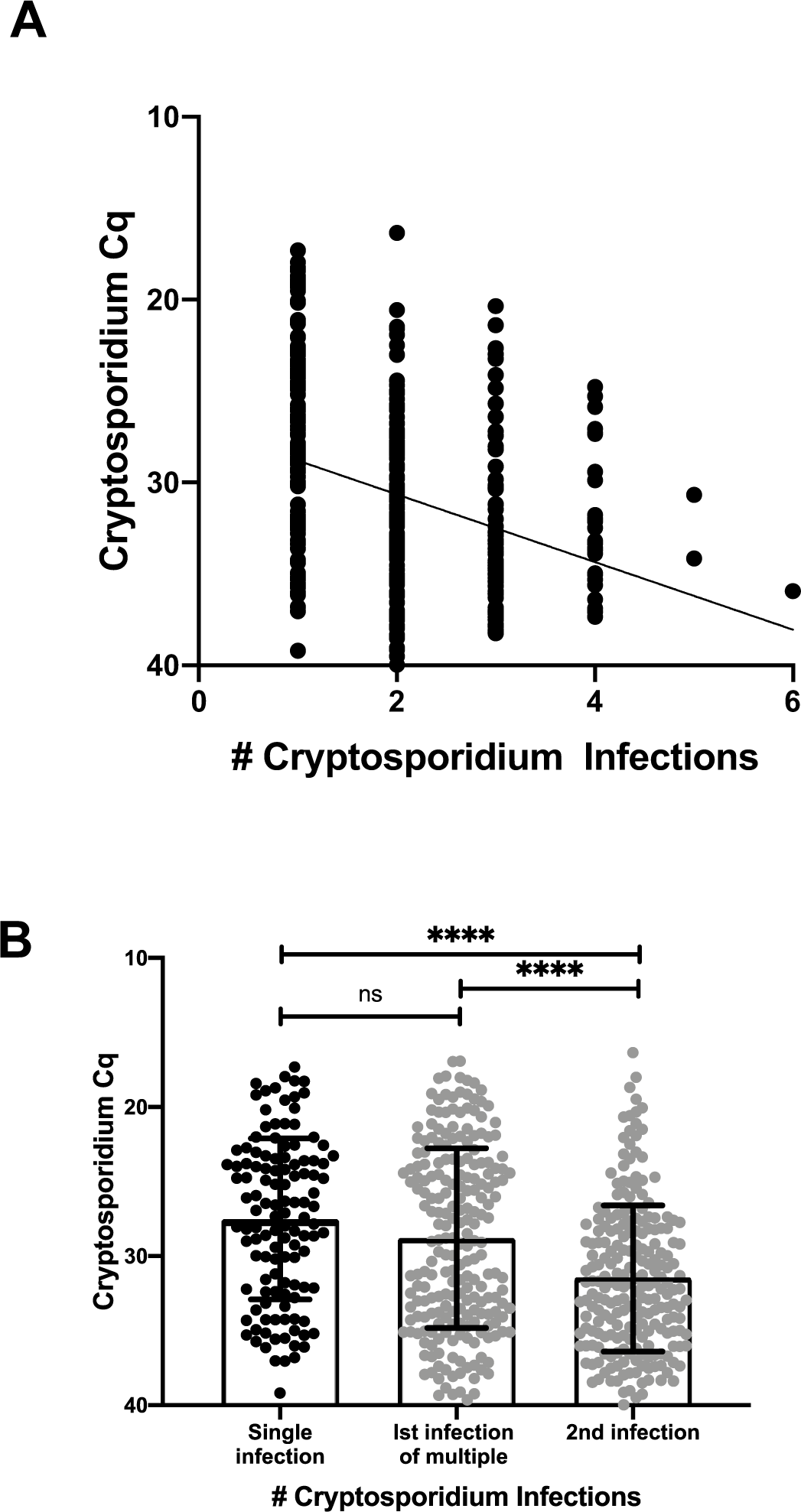
Parasite burden was lower in recurrent infections. A) Correlation between parasite burden and the number of *Cryptosporidium* infections. Each symbol represents the first detectable sample of an individual infection. Y-axis represents the quantitative cycle (Cq) of the diagnostic pan-Cryptosporidium PCR assay. X-axis shows the number of *Cryptosporidium* infections that had occurred in this child. The line represents the slope (−1.85 ± 0.21) and Y-intercept (26.95 ± 0.44) estimated from the GEE model with the exchangeable correlation structure (p<0.0001) B) Comparison of single infections (black symbol) with those that are part of a series (gray symbol). Bar graph (indicating data mean ± standard deviation) with individual data points. Each symbol on the box plot represents the first positive sample of an individual infection. X-axis refers to Infection number and, if the first infection, whether a second *Cryptosporidium* infection took place in the 3 years of life. Y-axis represents the quantitative cycle (Cq) of the diagnostic pan-Cryptosporidium PCR assay. Horizontal bars represent the result of a non-parametric Kruskal-Wallis test **** indicates p<0.0001

**Table 2.**
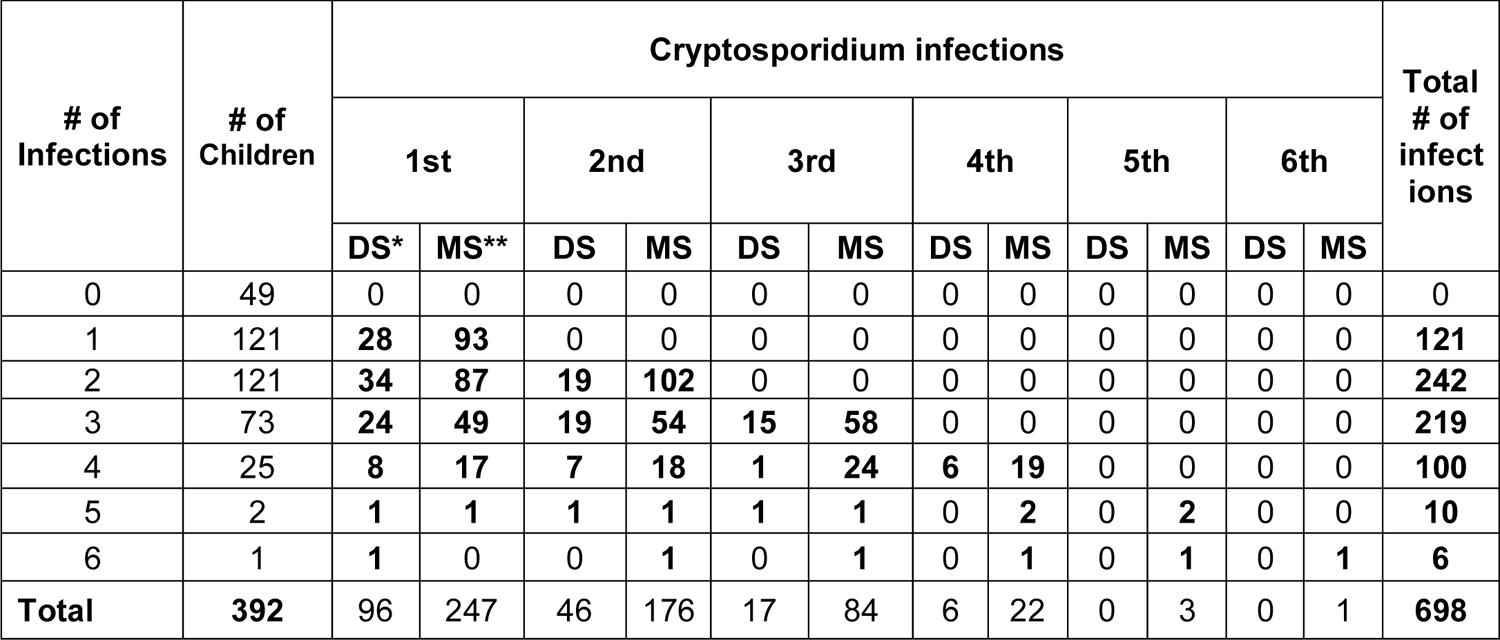
Distribution and Clinical Characterization of repeated *Cryptosporidium*

| # of Infections | # of Children | Cryptosporidium infections |  |  |  |  |  |  |  |  |  |  |  | Total # of infect ions |
| --- | --- | --- | --- | --- | --- | --- | --- | --- | --- | --- | --- | --- | --- | --- |
|  |  | 1st |  | 2nd |  | 3rd |  | 4th |  | 5th |  | 6th |  |  |
|  |  | DS* | MS** | DS | MS | DS | MS | DS | MS | DS | MS | DS | MS |  |
| 0 | 49 | 0 | 0 | 0 | 0 | 0 | 0 | 0 | 0 | 0 | 0 | 0 | 0 | 0 |
| 1 | 121 | 28 | 93 | 0 | 0 | 0 | 0 | 0 | 0 | 0 | 0 | 0 | 0 | 121 |
| 2 | 121 | 34 | 87 | 19 | 102 | 0 | 0 | 0 | 0 | 0 | 0 | 0 | 0 | 242 |
| 3 | 73 | 24 | 49 | 19 | 54 | 15 | 58 | 0 | 0 | 0 | 0 | 0 | 0 | 219 |
| 4 | 25 | 8 | 17 | 7 | 18 | 1 | 24 | 6 | 19 | 0 | 0 | 0 | 0 | 100 |
| 5 | 2 | 1 | 1 | 1 | 1 | 1 | 1 | 0 | 2 | 0 | 2 | 0 | 0 | 10 |
| 6 | 1 | 1 | 0 | 0 | 1 | 0 | 1 | 0 | 1 | 0 | 1 | 0 | 1 | 6 |
| Total | 392 | 96 | 247 | 46 | 176 | 17 | 84 | 6 | 22 | 0 | 3 | 0 | 1 | 698 |

To investigate if an initial high burden infection provided better protection against future infections with the *Cryptosporidium* parasite, we compared the Cq values of the infections in children who only had one infection in the first three years of life vs the Cq values of the first infection in children that had repeated infections (Fig 2B). The mean Cq values were similar in both cases (single infections: Cq 27.6 ± 5.4: 1^st^ infection of multiples: 28.8 ± 6.0) and significantly lower than that in subsequent second infections where infections were >1 (Cq second infection: 31.5 ± 4.9).

We next evaluated if the lower parasite burden in repeat infections was influenced by the age of the children. As most recurrent *Cryptosporidium* infections occurred in older children, (Table 1) we analyzed a subset of the *Cryptosporidium* positive samples corresponding to the first to fifth infections in children aged between 2.25 and 2.75 years. The parasite burden measured by qPCR remained significantly lower in the recurrent infections (Fig 3). The negative relationship of lower parasite burden with repeated infections was not an artefact of PCR inhibitors in the stool of older children because detection of the Phocine herpesvirus (PhHV) DNA included as internal extraction control [29] was not significantly affected by the number of prior *Cryptosporidium* infections. We concluded that repeat infections had a lower parasite burden.

**Fig 3.**
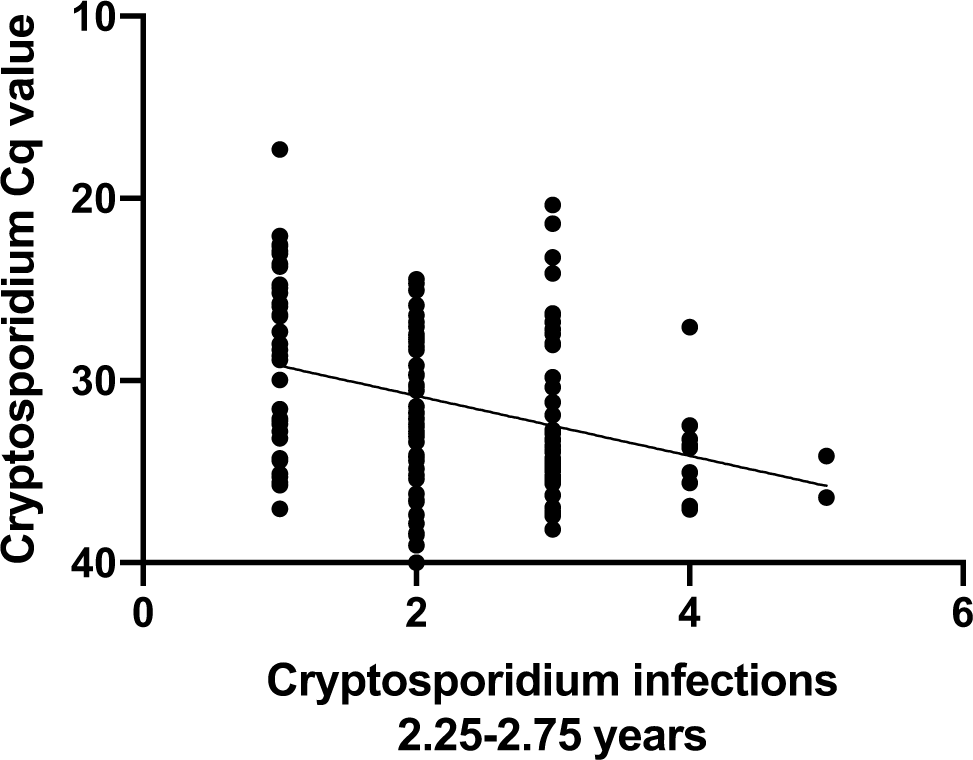
Parasite burden in older children

The amount of parasite in stool was determined as a function of the number of *Cryptosporidium* infections in a child by linear regression. The analysis was restricted to children between 2.25 and 2.75 years of age (n=140). Each symbol represents the first detectable sample of an individual infection. Y-axis, quantitative cycle of the diagnostic pan-Cryptosporidium PCR assay (Cq). X-axis, the number *Cryptosporidium* infections that had occurred in each child. Slope: −1.65, R squared value: 0.01125, Significance p<0.0001.

The duration of diarrheal disease was similar in the first infection and later reinfections (single infection: 4.8 ± 2.9 days; primary infection: 5.5 ± 3.9 days; later infections: 5.2 ± 3.6 days), however, the proportion of the diarrhea-associated *Cryptosporidium* infections decreased in the recurrent infections (Chi-squared test for trend p=0.011) (Table 1). We concluded that the repeated *Cryptosporidium* infections were more likely to be sub-clinical.

### *Cryptosporidium* and growth faltering

Growth faltering (low height for age; HAZ score) was analyzed from the 3 year old children based on the number of *Cryptosporidium spp.* infections [0 - 3 years] (both diarrhea and sub-clinical) (Table 2, Fig S5 & 6). The association between cryptosporidiosis and HAZ score at three years was examined by multiple regression in order to account for the effect of the confounding variables previously identified (Table 3) [13, 21].

**Table 3.**
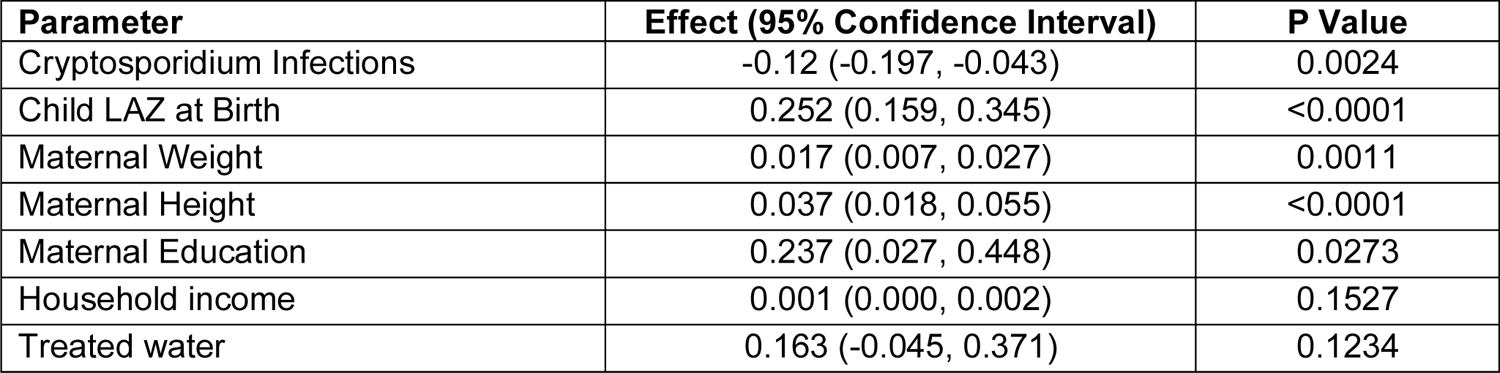
Regression Analysis using selected predictors to test the association of Cryptosporidiosis with Height-for-Age Scores at 3 Years

*Cryptosporidium* infection was negatively associated with the HAZ score at 3 years after adjusting for birth length-for-age (LAZ) score and maternal weight and education: each *Cryptosporidium* infection reoccurrence resulted in a decrease in HAZ score (Δ 0.12) at 3 years (Table 5). No significant relation was found between malnutrition at birth (LAZ score) and total number of *Cryptosporidium* infections during the follow-up (Fig 4A). However, the total number of *Cryptosporidium* infections was negatively associated with HAZ score at 3 years (Fig 4B, regression coef=-0.152, p=0.0004) The association between HAZ and *Cryptosporidium spp*. infections was unaffected by whether the event was a sub-clinical infection or diarrheal disease (Fig S7).

**Fig 4.**
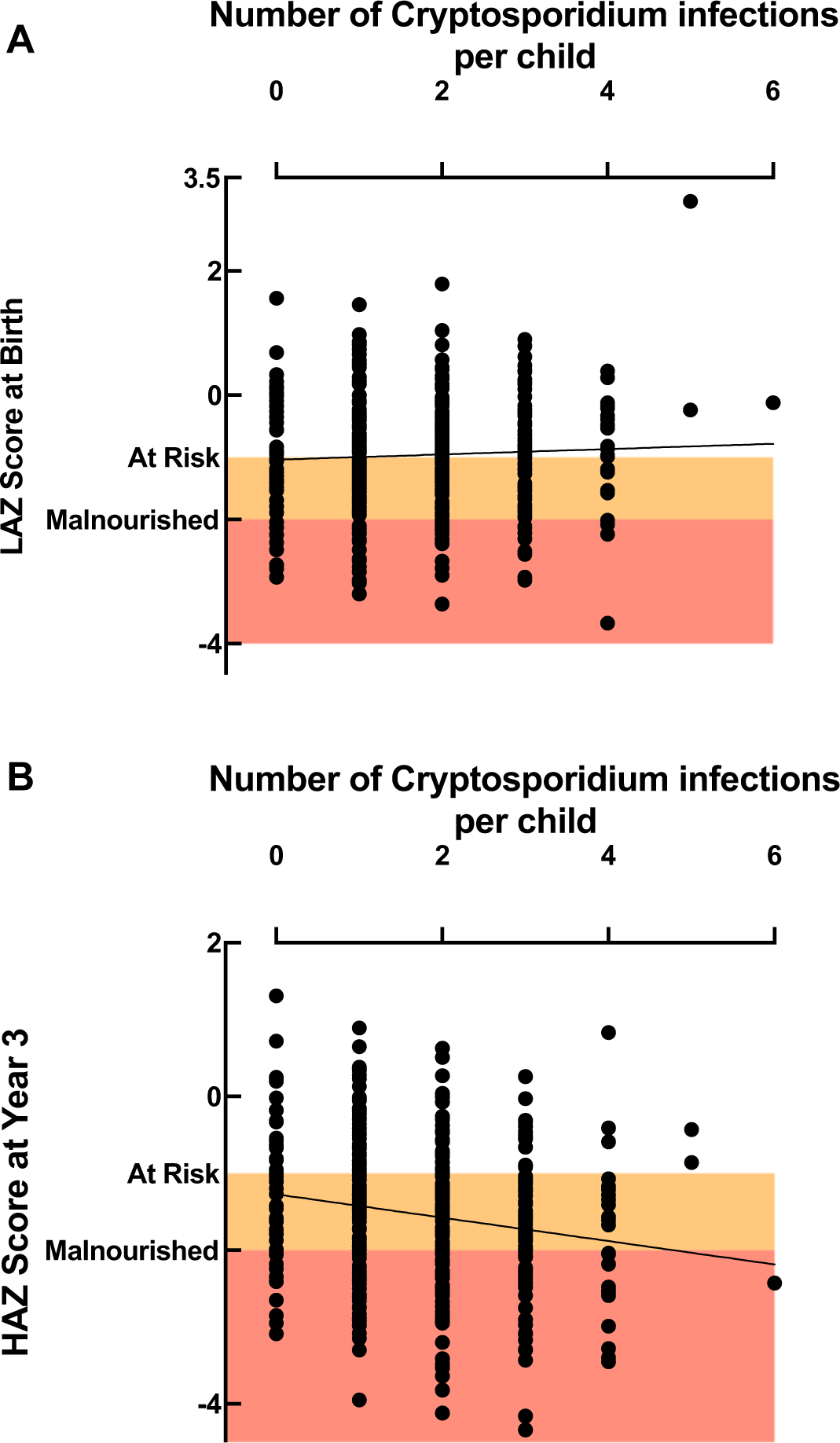
Cryptosporidiosis frequency was associated with growth faltering distinct from the impact of birth nutritional status A) Relationship between length for age z score (LAZ) at birth (Y-axis) and the total number of *Cryptosporidium spp.* infections (X-axis). The slope was not significantly different from one. B) Relationship between height for age z score (HAZ) at 3 years (Y-axis) and the total number of *Cryptosporidium spp.* infections (X-axis). Slope: −0.152 ± 0.0429, R squared value: 0.0313, Significance p=0.0004. Children were defined to be at risk for growth faltering with a LAZ or HAZ score <-1 and malnourished at LAZ or HAZ score <-2. Orange box: birth LAZ or 3 year HAZ score −1 to −2; red box: birth LAZ or 3 year HAZ or score < −2.

**Table 5.**
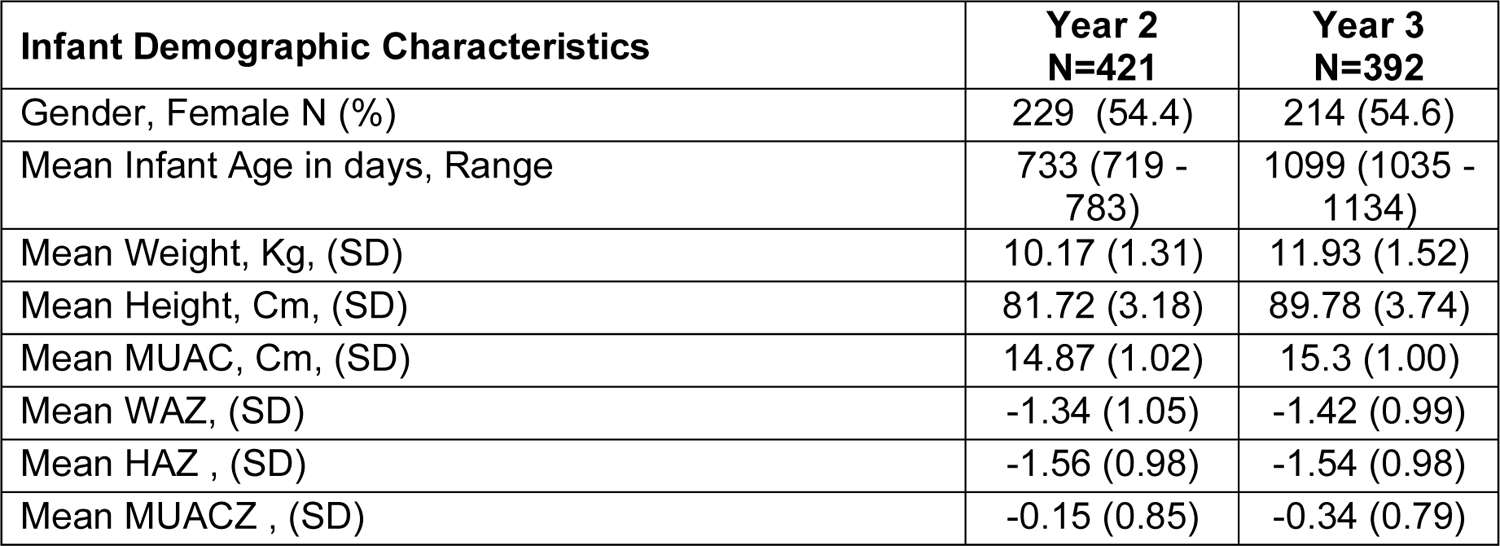
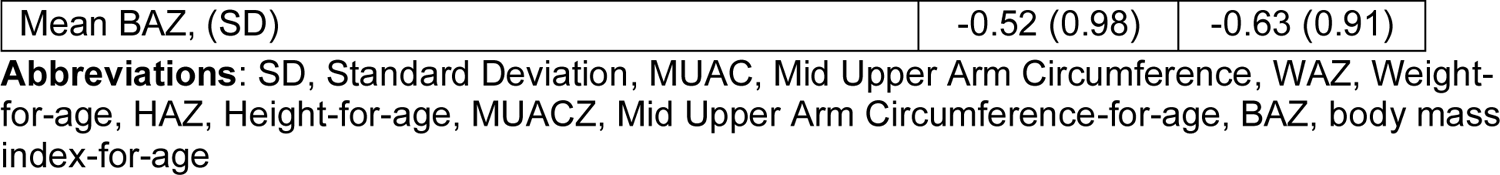
Infant demographic characteristics.

| Infant Demographic Characteristics | Year 2<br>N=421 | Year 3<br>N=392 |
| --- | --- | --- |
| Gender, Female N (%) | 229 (54.4) | 214 (54.6) |
| Mean Infant Age in days, Range | 733 (719 - 783) | 1099 (1035 - 1134) |
| Mean Weight, Kg, (SD) | 10.17 (1.31) | 11.93 (1.52) |
| Mean Height, Cm, (SD) | 81.72 (3.18) | 89.78 (3.74) |
| Mean MUAC, Cm, (SD) | 14.87 (1.02) | 15.3 (1.00) |
| Mean WAZ, (SD) | -1.34 (1.05) | -1.42 (0.99) |
| Mean HAZ, (SD) | -1.56 (0.98) | -1.54 (0.98) |
| Mean MUACZ, (SD) | -0.15 (0.85) | -0.34 (0.79) |
| Mean BAZ, (SD) | -0.52 (0.98) | -0.63 (0.91) |

**Abbreviations:** SD, Standard Deviation, MUAC, Mid Upper Arm Circumference, WAZ, Weight-for-age, HAZ, Height-for-age, MUACZ, Mid Upper Arm Circumference-for-age, BAZ, body mass index-for-age

Other measurements that are used as indicators of malnutrition were also significantly associated with the number of *Cryptosporidium* infections. These included mid-upper arm circumference (MUAC) (Fig S8A; MUACZ vs. number of *Cryptosporidium* infections slope: −0.088; p=0.0123 and weight-for-age (WAZ score) (Fig S8B) (linear regression analysis slope: −0.115 p=0.0093). However, neither BAZ (body-mass-for-age) (Fig S8C), used to measure acute protein-energy malnutrition or wasting (WHZ) were affected by a history of *Cryptosporidium* infections (Fig S8D).

The Pearson correlations among the number of *Cryptosporidium* infections, LAZ at birth, diarrheal episodes and HAZ at year 3 are shown in Fig 5A (*Cryptosporidium* infections: HAZ at year 3: coef = −0.18, p=0.024; *Cryptosporidium* infections: diarrheal episodes captured (all causes): coef = 0.22, p>0.0001; HAZ at year 3: LAZ at birth: coef = 0.28, p= 0.008). As expected, a significant correlation existed between LAZ at birth and HAZ at year 3 (simple linear regression p<0.0001 Fig 5B).

**Fig 5.**
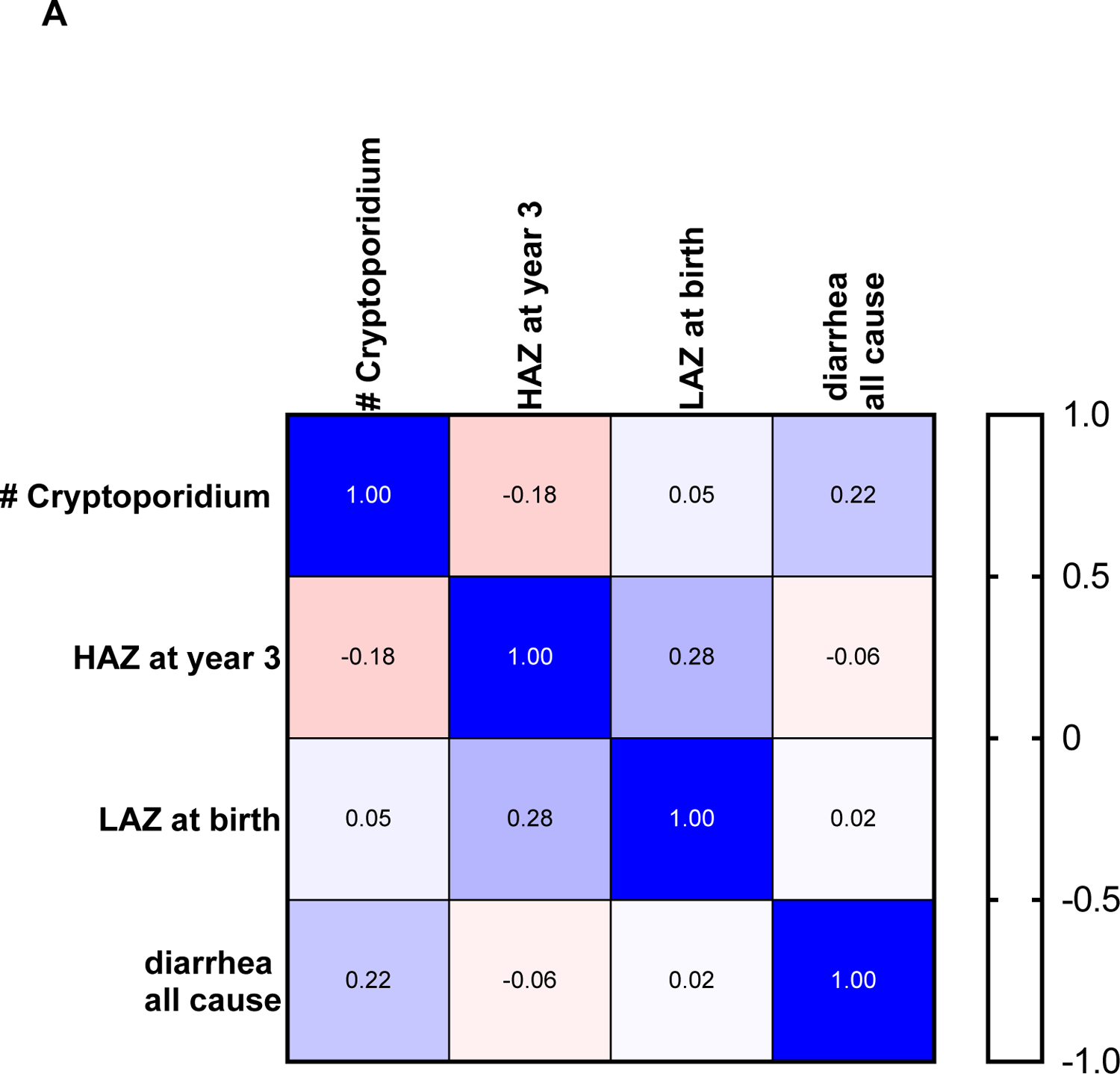
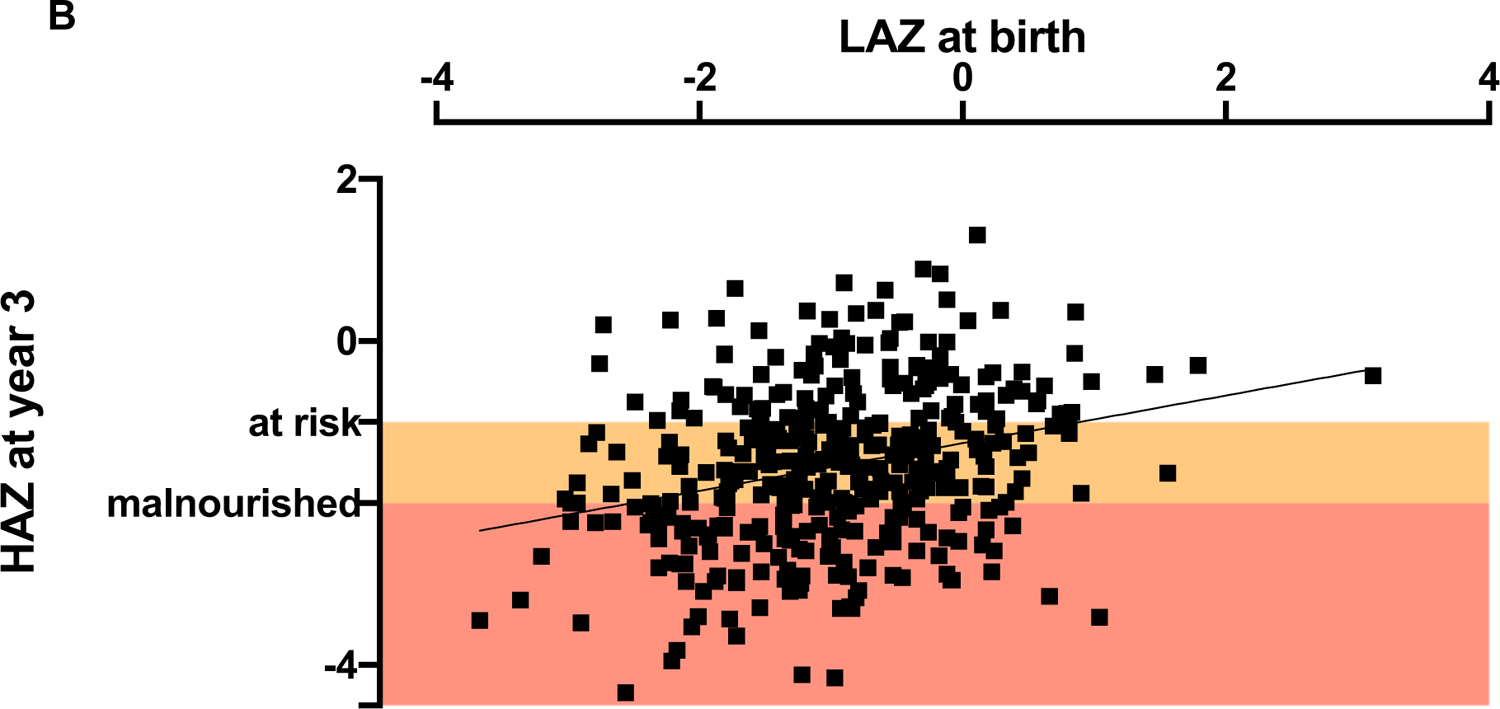
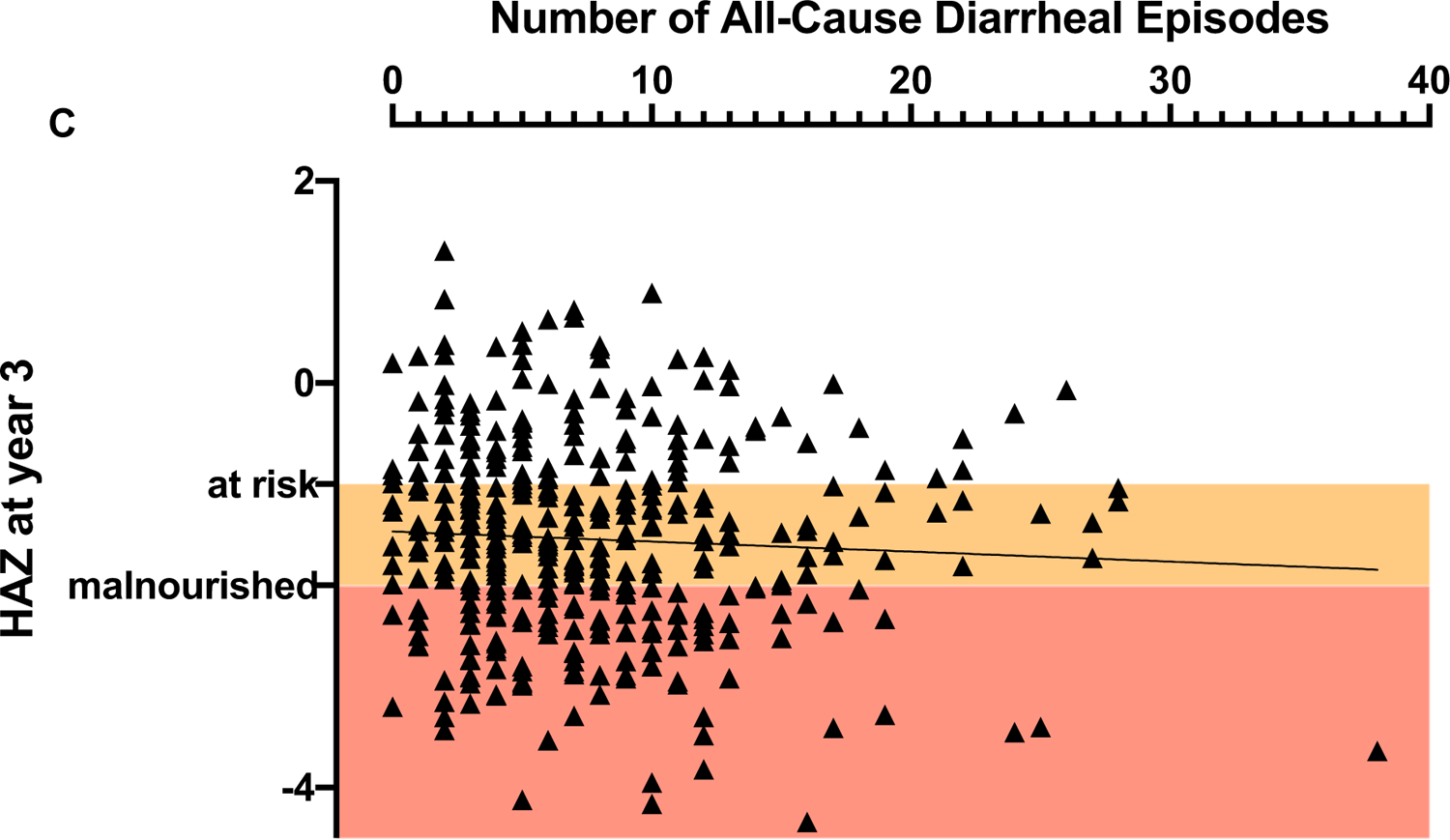
Correlates of cryptosporidiosis-associated growth-faltering. A) Correlation matrix of cryptosporidiosis, all-cause diarrhea, LAZ at birth and HAZ at 3 years, calculated using Pearson r. Bar on the right indicates strength and direction of association. B) Comparison of three year-HAZ with birth LAZ. (Slope: −0.294 ± 0.05; R squared value: 0.08; Significance p < 0.0001). C) Relationship of all-cause diarrhea with HAZ at 3 years of age (p = NS).

Enteric pathogens are endemic in the Bangladesh study population [30] and as a consequence, infants enrolled in the study cohort had repeated diarrheal episodes of which only some were associated with infection with the *Cryptosporidium* parasite. However, while *Cryptosporidium* infections (diarrheal and sub-clinical) were significantly associated with child HAZ at year 3 (Pearson’s correlation p=0.0004), the number of all-cause diarrheal episodes was not (Fig 5C; S2 Table 3). This result supported our conclusion that this growth shortfall was specifically associated with recurrent cryptosporidiosis.

### Mucosal IgA against the sporozoite Cp23 protein was associated with protection from growth faltering

In previous work it was shown in this cohort that a high level (> mean value) of fecal anti-Cp23 IgA at one year of age was associated with an increased resistance to cryptosporidiosis through age three [14, 20]. Here we additionally discovered that children with high levels (upper 50^th^ percentile) of fecal anti-Cp23 IgA at one year of age were protected from growth faltering through year 3 (Fig 6). Subgrouping the children into Group 2a (never infected by evidence of anti-Cp23 IgA levels and diagnostic qPCR assays; n=20) and Group 2b (diagnostic qPCR positive only; n=185) versus Group 1 children with high levels of IgA at one year (n=171) did not alter the association with growth faltering (Fig S9A). Analysis of the fecal IgA antibodies against a second sporozoite peptide (Cp17) was also performed. Although a similar trend was observed the difference in year 3 HAZ was not significantly different (Fig S9B). A high level of fecal anti-Cp23 at one year was not associated with any drop in the parasite burden at the next C*ryptosporidium* infection (first subsequent new infection: Cq of IgA high responders 28.2 ± 5.6 versus 27.6 ± 5.6 p=0.43).

**Fig 6.**
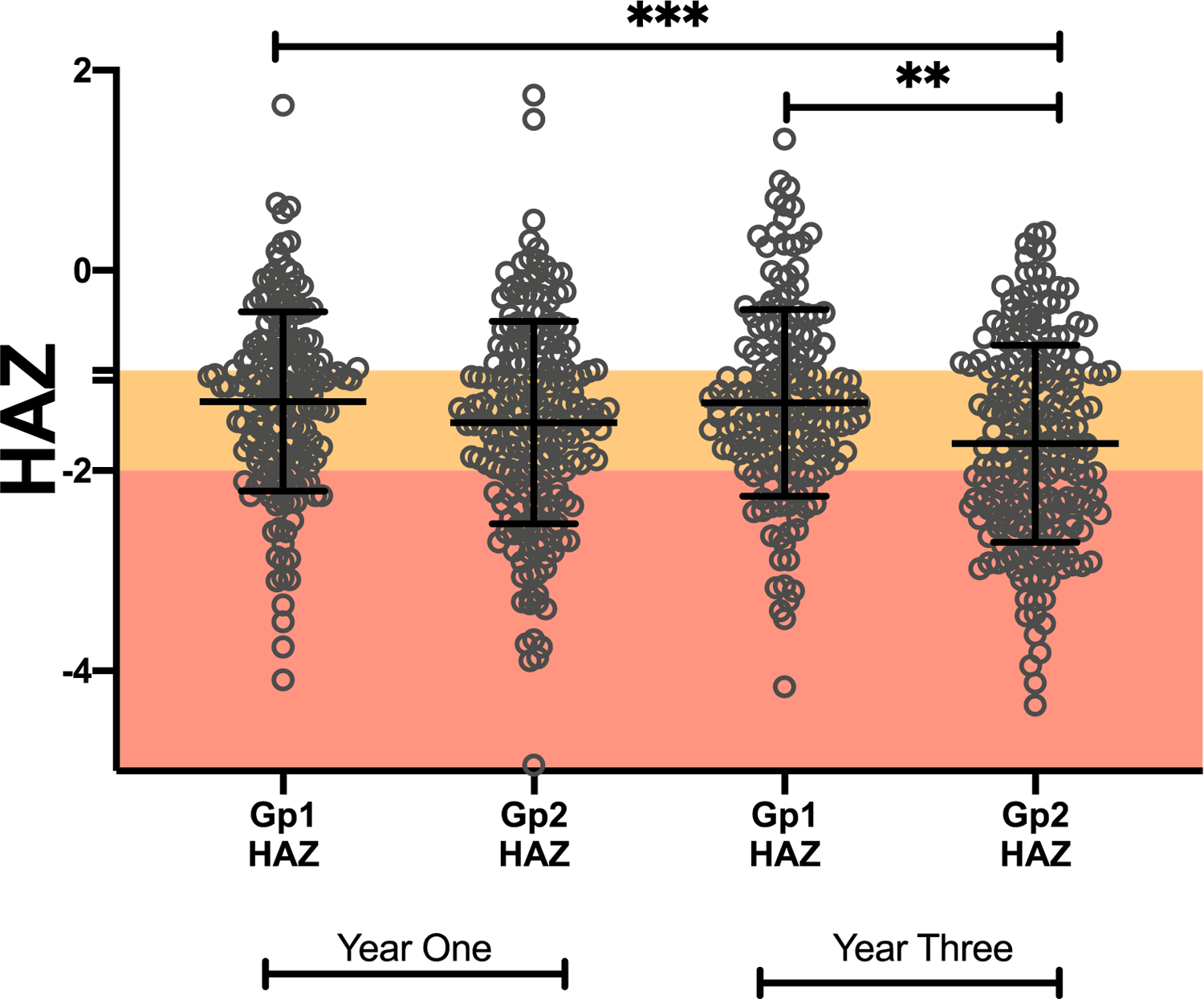
High anti-Cp23 IgA levels were associated with a reduction in cryptosporidiosis-associated growth-faltering.

Group 1 and 2 children were in the upper and lower 50th percentile for fecal IgA anti-Cp23 respectively. HAZ are shown for children in both year one and year three of life. Mean ± standard deviation with individual data points. Horizontal bars represent the result of a non-parametric Kruskal-Wallis test ***p<0.001, **p<0.01

## Discussion

The key finding of this paper is that naturally acquired immunity protects from *Cryptosporidium* diarrhea but does not provide sterilizing immunity. The importance of this observation is two-fold: first it indicates that transmission likely occurs in semi-immune populations; and second that continued sub-clinical infections increase the risk of infection-related growth faltering. Encouragingly however, acquired immunity associated with high levels of mucosal IgA against the Cp23 cryptosporidium sporozoite antigen were associated with protection from malnutrition.

Many previous studies on cryptosporidiosis have focused on the health impact of diarrhea-associated cryptosporidiosis [7,12,31–34]. However sub-clinical disease, as opposed to infection accompanied by diarrhea, may also have long term effects on child health. The link between sub-clinical cryptosporidiosis and malnutrition is now well known if not yet well understood [8,9,13,32]. In a recent study the global prevalence of cryptosporidiosis in people without diarrheal symptoms was 4.4% (95% confidence interval 2.9 - 6.3)[35]. During the 3 years of this study 212 children (54%) had only sub-clinical *Cryptosporidium* infections. This longitudinal study allowed us to take an in depth look at the role of sub-clinical reinfections in the exacerbation of growth faltering [13,22,25].

Anthropometric measurements are reliable non-invasive methods to monitor child malnutrition. The most commonly used metrics are a shortfall in child growth (low height for age: HAZ score) a consequence of chronic undernutrition and wasting (exemplified by a low weight for height: WHZ score). In line with most studies our results show that a history of cryptosporidiosis was associated with a decrease in the HAZ score of children irrespective of infection severity [8,26,36,37]. Here we found that child growth was negatively impacted not only by the first episode of cryptosporidiosis, but both occurred and remained constant in succeeding infections, even though parasite burden and diarrheal disease decreased. This study has, therefore, shown that naturally acquired partial immunity was not effective at preventing growth faltering and that a control strategy focused on only preventing diarrheal cryptosporidiosis may not prevent the stunted growth associated with cryptosporidiosis.

A limitation of the current study is that it was not possible to unambiguously attribute an episode of diarrhea to *Cryptosporidium* because children in this community were infected with multiple enteropathogens at the same time [26, 30]. To mitigate the problem of correctly identifying *Cryptosporidium*-associated diarrheal infections these were defined as an episode of diarrhea accompanied by a new *Cryptosporidium* infection (i.e. the immediately preceding surveillance or diarrheal stool sample was negative for *Cryptosporidium*) [13]. A second limitation was that surveillance stool samples were collected at only monthly intervals which likely missed some subclinical infections, potentially underestimating the impact of cryptosporidiosis on child growth. The study however had notable strengths including most importantly its longitudinal design that combined collection of surveillance and clinical specimens with studies on child growth faltering.

The association of mucosal immunity to Cp23 with protection from growth faltering offers hope that a cryptosporidiosis vaccine could have a measurable impact on child health, even in the absence of absolute protection from infection.

## Acknowledgement

We are grateful for the participation of the parents and children in this study as well as the staff of the Emerging Infectious Diseases Division of icddr,b for contributing to this research.

## Author Contributions

CG, MK, RH and WP designed, and RH, WP, MA, JAW, CG and MK drafted the study protocols and WP directed the work described in this paper. MK, TA, BH, RT and AK acquired the data used in this manuscript. UN curated the study data. UN, JM, MK and CG analyzed the study data. CG wrote the first draft of the manuscript that was reviewed and amended by all the authors who also approved the final manuscript.

## Methods

### Child cohort

A total of 500 children were enrolled within one week of birth in an urban slum of Dhaka, Bangladesh beginning in June 2014 through March 2016 and were monitored for diarrheal diseases through bi-weekly home visits by trained field investigators. A monthly stool sample was also collected to evaluate asymptomatic infection and growth was measured every 3 months (“Cryptosporidiosis and Enteropathogens in Bangladesh”; ClinicalTrials.gov identifier NCT02764918). This area (Section 11 of Mirpur Thana) is densely populated with participants in this study having an average of 5.5 people living in 1.6 rooms. Annual median household income of participants was 14,000 Taka or approximately US $164 (Table 4). Anthropometric data was collected as previously described [13]. Each child was weighed on an electronic scale (kilograms, measured with electronic scale; TANITA, HD-314). Child height or length (depending on age) and mid-upper arm circumference were measured to the nearest 0.1 cm using a measuring board and plastic tape (Table 5). The height-for-age z score (HAZ); weight for age z score (WAZ); weight for height (WHZ); body mass index for age (BAZ); and mid-upper arm circumference for age (MUACZ) were calculated using the World Health Organization Anthro software (version 3.2.2) [13]. Children who had a HAZ score <-1 were defined as ‘at risk for malnutrition’ and HAZ < −2 as malnourished [27, 28]. Diarrhea was defined as ≥3 loose stools within a 24-hour period as reported by the child’s caregiver with episodes separated by a gap of at least 3 days. This paper reports the data from 392 infants who were followed through three years of age.

**Table 4.**
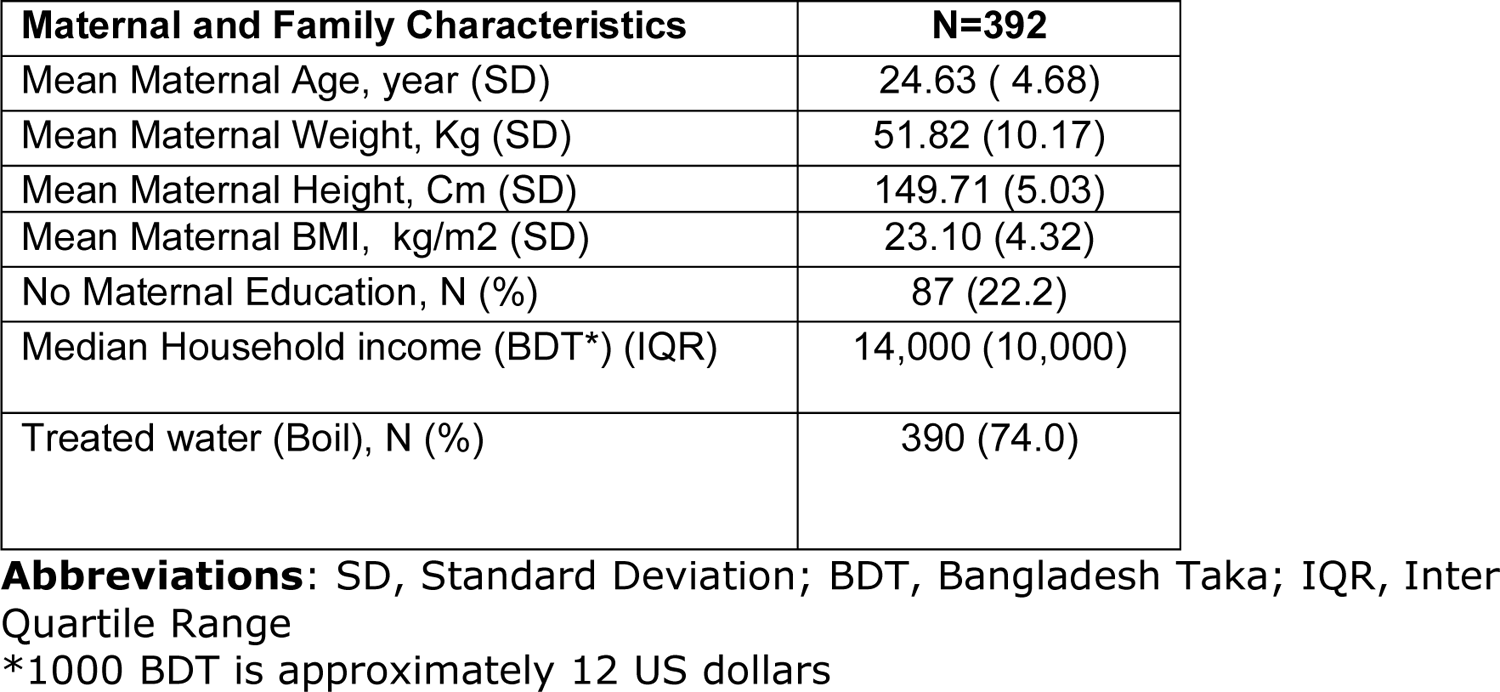
Maternal and family demographics.

| Maternal and Family Characteristics | N=392 |
| --- | --- |
| Mean Maternal Age, year (SD) | 24.63 ( 4.68) |
| Mean Maternal Weight, Kg (SD) | 51.82 (10.17) |
| Mean Maternal Height, Cm (SD) | 149.71 (5.03) |
| Mean Maternal BMI, kg/m<sup>2</sup> (SD) | 23.10 (4.32) |
| No Maternal Education, N (%) | 87 (22.2) |
| Median Household income (BDT*) (IQR) | 14,000 (10,000) |
| Treated water (Boil), N (%) | 390 (74.0) |

**Abbreviations:** SD, Standard Deviation; BDT, Bangladesh Taka; IQR, Inter Quartile Range
\*1000 BDT is approximately 12 US dollars

### Ethics Statement

The study was approved by the Ethical and Research Review Committees of the International Centre for Diarrhoeal Disease Research, Bangladesh (PR-13092) and by the Institutional Review Board of the University of Virginia (IRB#20388). Informed written consent was obtained from the parents or guardians for the participation of the subjects in the study.

### Sampling and specimen testing

Fresh stool samples collected in the field were placed on ice and then brought to the lab on the same day and frozen within 6 h of collection (Fig S1). Stool specimens were collected from children every month (monthly surveillance) and during episodes of diarrhea. A modified Qiagen stool DNA extraction protocol with 95°C incubation and a 3-minutes bead–beating step was used to extract DNA [13] (Fig S2). These samples were tested with a multiplex qPCR assay which utilizes pan-Cryptosporidium primers and probes targeting the 18S rDNA gene and primers and probes to detect the Phocine herpesvirus (PhHV) extraction control (obtained from the European Virus Archive Global organization) as previously described (Fig S3). All samples with a cycle threshold of ≤ 40 for cryptosporidium were used in this analysis [13]. In year 3 the diagnostic qPCR assay was not able to be completed on 0.9% of the collected diarrheal and 4.6% of the monthly surveillance samples (Table S1).

Infection with *Cryptosporidium* was defined as detection of *Cryptosporidium* DNA by qPCR from stool. PCR-positive samples were classified as a separate infection if occurring greater than 65 days after the preceding positive sample [13]. The *Cryptosporidium* infection phenotype (diarrheal or sub-clinical) was based upon symptoms at the time of detection of the first *Cryptosporidium* - positive stool sample, whether diarrheal stool or monthly surveillance.

### Statistical analysis

Descriptive statistics were expressed in mean ± standard deviation for continuous variables and as frequencies and proportions for categorical variables. The frequency of repeated *Cryptosporidium* infections in the first 3 years of life was summarized for diarrhea and sub-clinical infections separately and their differences were evaluated with the χ^2^test. To account for within-child correlations among repeated *Cryptosporidium* infections, the relationship between parasite burden and the number of repeated *Cryptosporidium* infections was evaluated using the Generalized Estimating Equation (GEE) for repeated measurements, assuming an exchangeable correlation structure. Pearson correlation was calculated for univariate association of individual predictors with HAZ at 3 years. Since confounders such as LAZ at birth, maternal weight and height, maternal education, household income and access to treated water were previously shown to impact HAZ [8, 13], a multivariable linear regression was performed to evaluate the association between *Cryptosporidium* infection and HAZ at 3 years after adjusting for these factors (Table 3). Similarly, a multiple regression analysis was performed to independently evaluate whether the number of episodes of diarrhea, irrespective of the causative pathogen, was associated with HAZ at 3 years. Analyses were performed using both the GraphPad Prism version 8.4.3 for Mac, (GraphPad Software, San Diego, California USA,), SAS 9.4 (Raleigh, NC) and R version 3.3.3, 32-bit.

## Supplemental Figures and Tables

**Fig S1.**
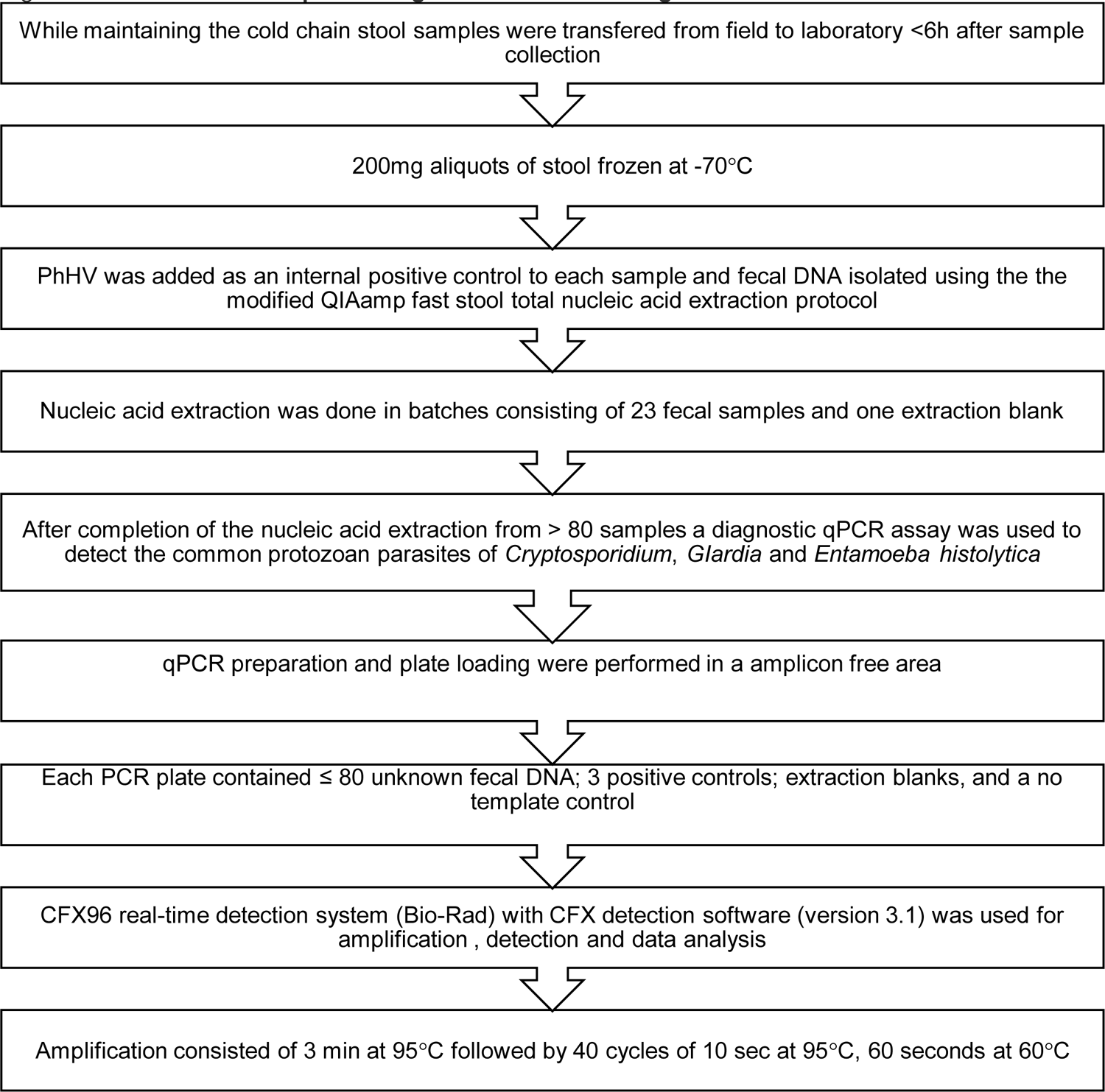
Flow chart of stool processing and molecular testing

**Fig S2.**
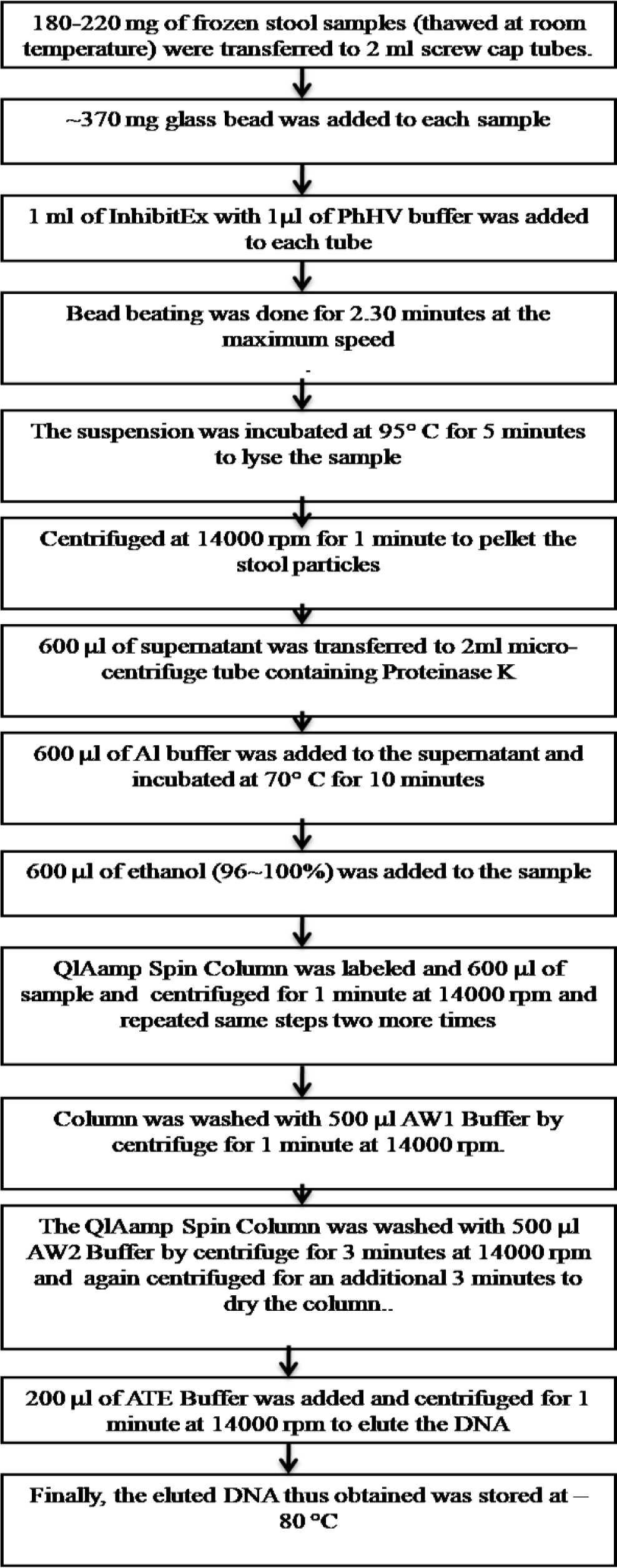
**Flow chart of stool TNA extraction procedure using QIAamp Fast DNA Stool Mini Kit from fresh or frozen stool samples.**

**Fig S3.**
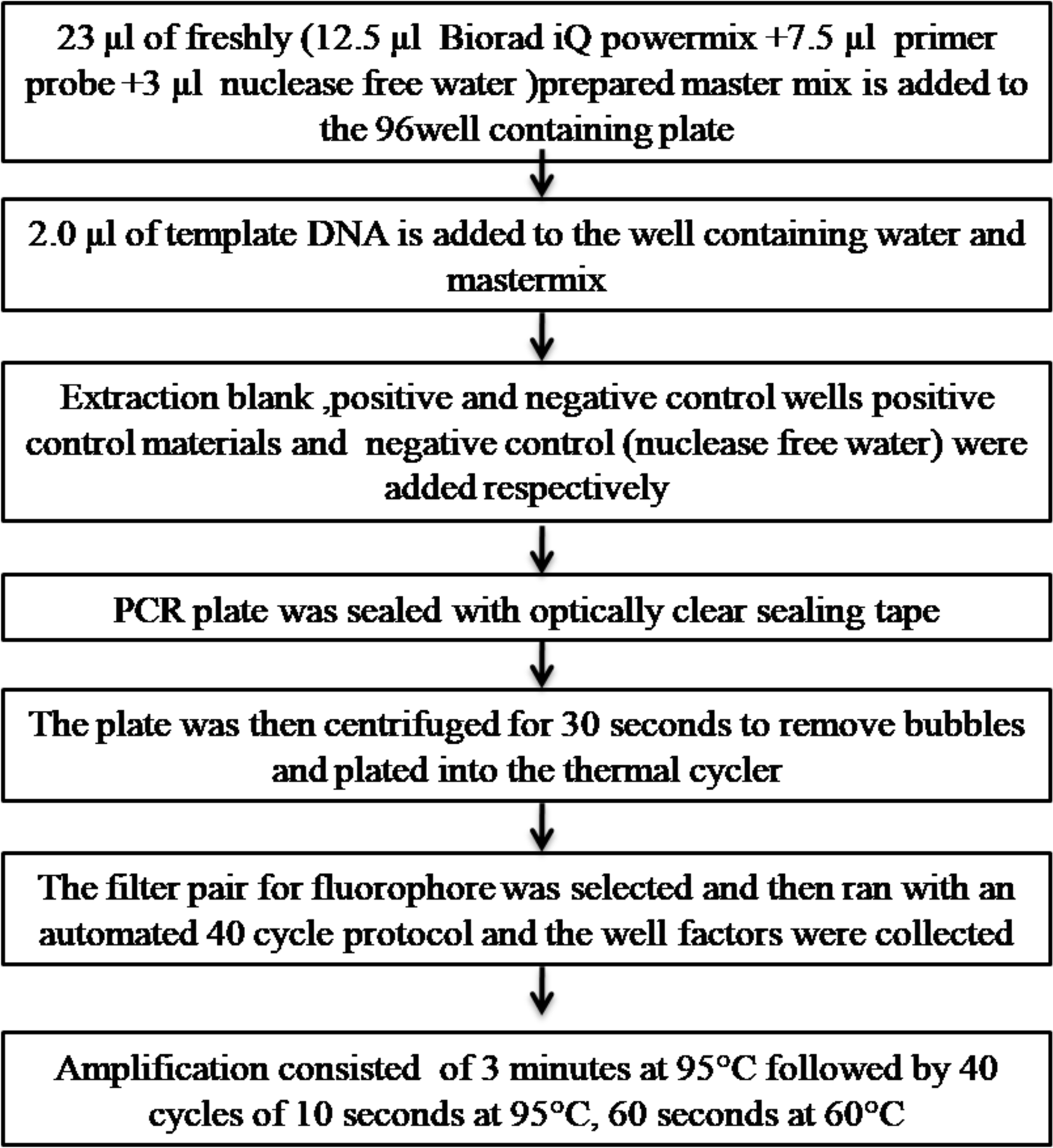
**Flow chart of Multiplex qPCR of Cryptosporidium, Giardia, Entameoba histolytica by targeting the18S gene**

**Fig S4.**
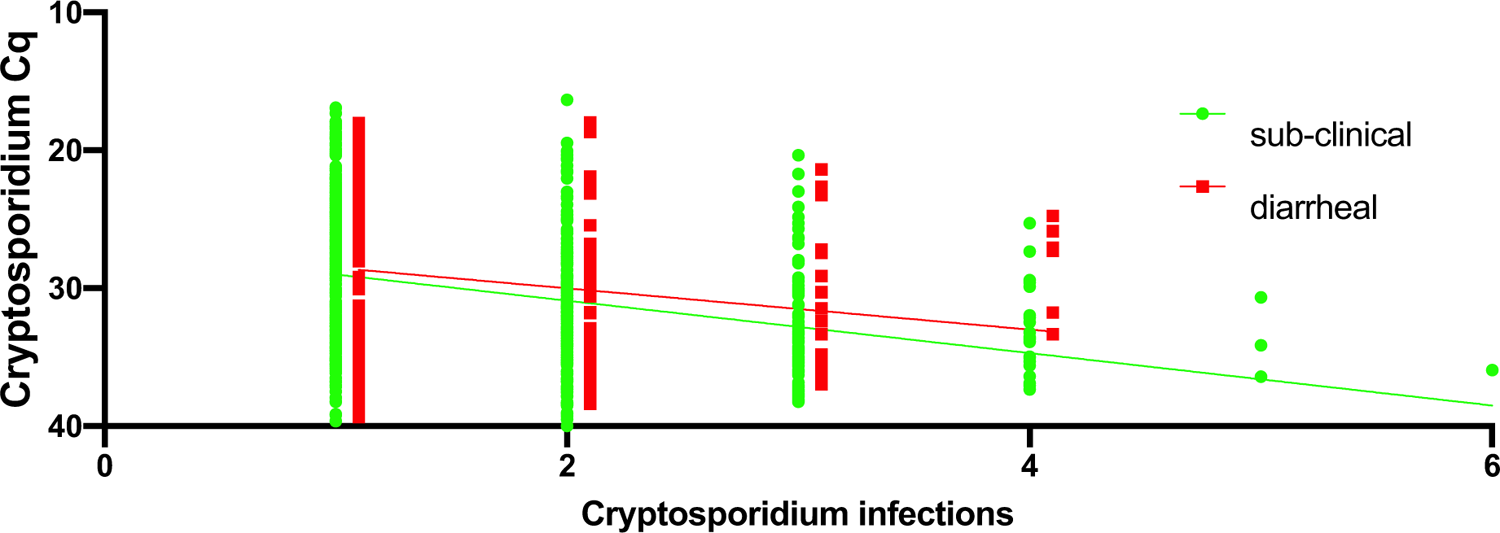
Parasite Burden in diarrheal and sub-clinical infections. Relationship between Parasite Burden and the number of recurrent *Cryptosporidium* infections. Each symbol represents the first detectable sample of an individual infection. Y-axis, quantitative cycle of the diagnostic pan-Cryptosporidium PCR assay (Cq). X-axis, the total number of *Cryptosporidium* infections. The infection was designated as either diarrheal (red) or sub-clinical (green) based on the current infection phenotype. The data from diarrheal cases was offset to improve data visualization. To account for within-child correlations among repeated *Cryptosporidium* infections, the generalized estimating equation (GEE) method for repeated measurements were used with exchangeable correlation structure. As the intercept of the diarrheal and sub-clinical models was not statistically different the common intercept (27.02 ± 0.45) was used. The slope of the data derived from the sub-clinical (1.9 ± 0.2) and diarrheal (1.49 ± 0.31) exchangeable models were not significantly different from each other (p=0.071) although both were statistically different from zero (p<0.0001).

**Fig S5.**
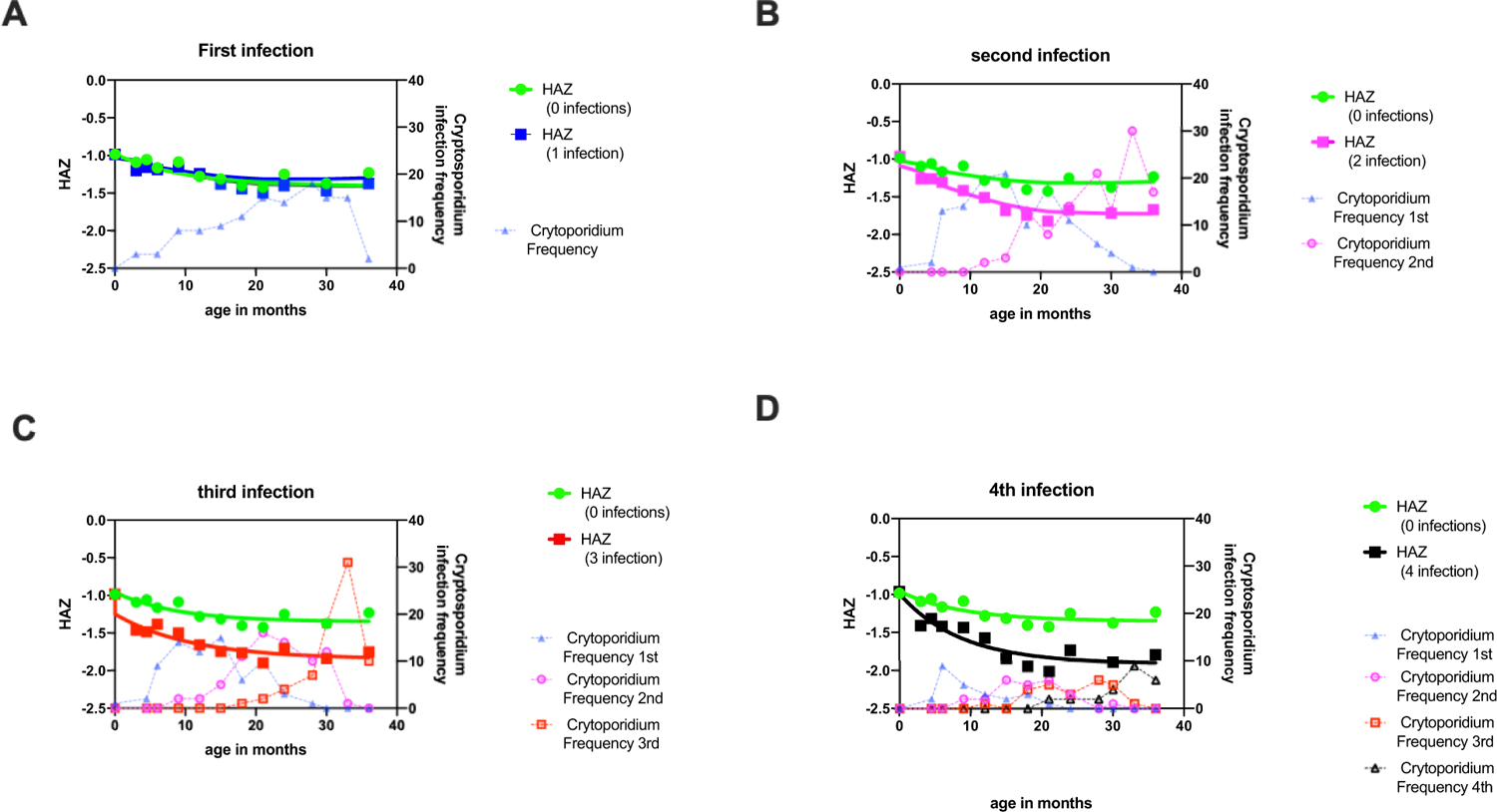
Distribution of repeated *Cryptosporidium* infections x axis child age in months; left y-axis child HAZ scores; right y-axis frequency of *Cryptosporidium* (diarrheal and sub-clinical) infections (shown as the number that occurred per the age of the child in months). All graphs include as a reference the HAZ score of children where no *Cryptosporidium* infections were detected (green circle and line). *Cryptosporidium* infections: Light blue triangle dotted blue connection line: infection one; purple circle and dotted line: infection two; light red square and dotted line: infection three; black triangle and dotted line: infection four A) blue symbol and solid line HAZ score of children who had one *Cryptosporidium* infections by 3 years of age B) purple square and solid line HAZ score of children who had two *Cryptosporidium* infections by three years of age C) red square and solid line HAZ score of children who had three *Cryptosporidium* infections by three years of age D) black square and solid line HAZ score of children who had four *Cryptosporidium* infections by three years of age

**Fig S6.**
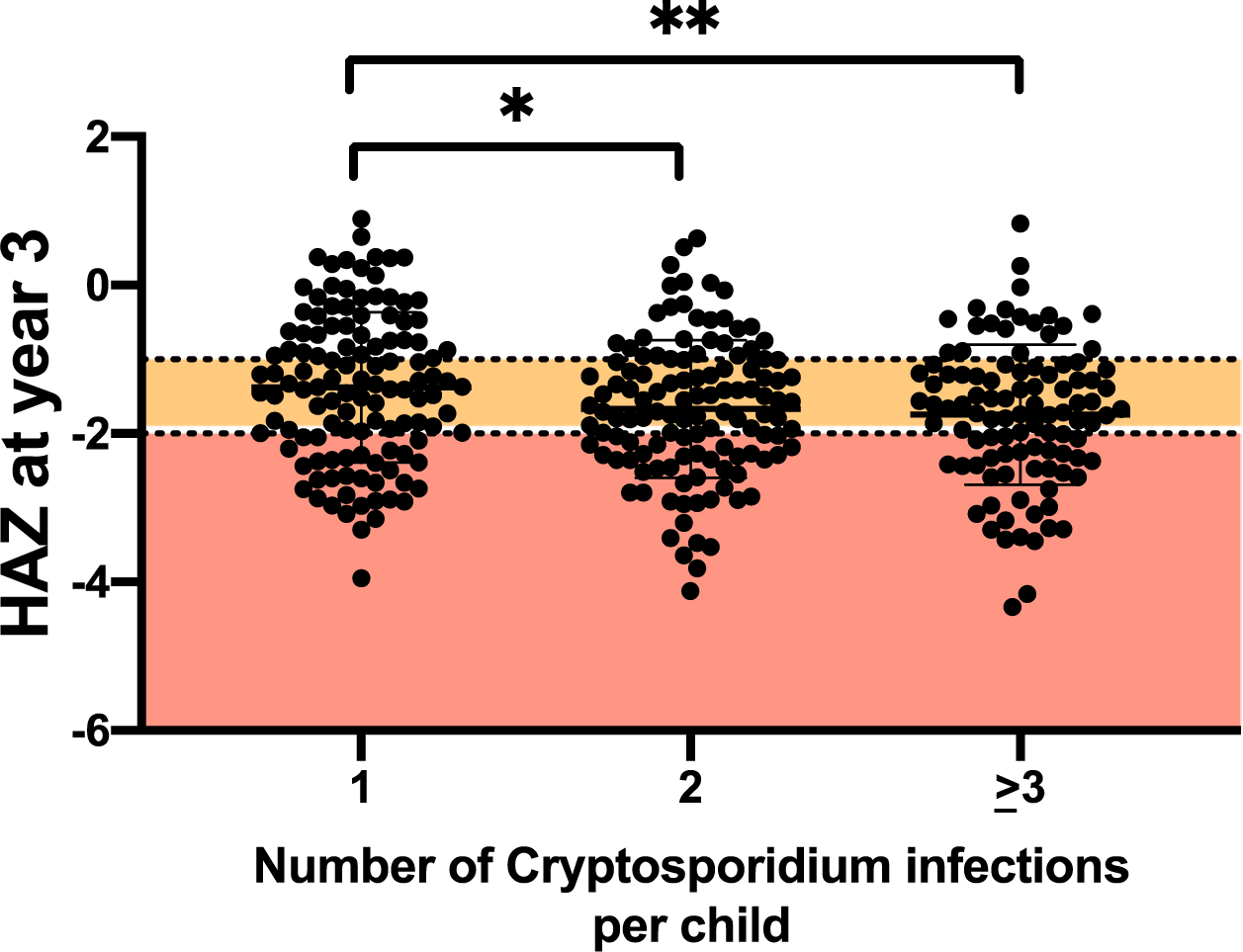
Recurrent cryptosporidiosis results in greater growth faltering. Each symbol represents a single child. Box plot comparing the height for age z score at 3 years (HAZ) (Y-axis) mean and standard deviation shown Children were considered to be at Risk for malnutrition is they have a HAZ score <-1 and malnourished at HAZ-2: orange box: 3-year HAZ score −1 to −2; red box 3-year HAZ score < −2. X-axis Number of Cryptosporidium infections. Bar indicates the result of a non-parametric Kruskal-Wallis test for multiple comparisons * indicates p<0.05 ** indicates p<0.01

**Fig S7.**
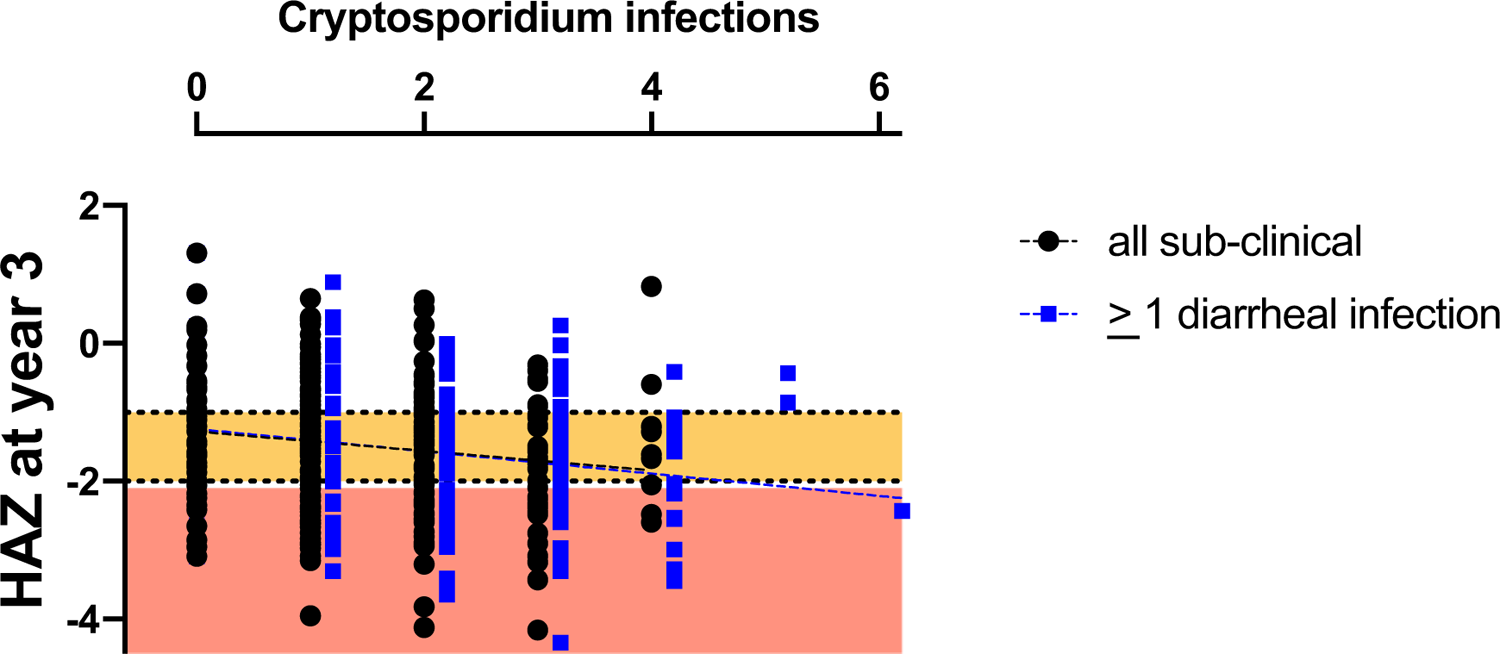
Comparison of Cryptosporidiosis associated Growth-faltering in diarrheal and sub-clinical infections. Graphs show results from a simple linear regression with each symbol representing a single child. Black symbols represent children who were never infected or had sub-clinical infections. The blue symbols indicate children who have had one or more than one episodes of diarrhea-associated cryptosporidiosis. Height for age (HAZ) z score at 3 years is shown on the Y-axis. The slope of the diarrheal-associated and sub-clinical groups are identical. Pooled Slope: −0.1545. Children are considered to be at Risk for malnutrition if they have a HAZ score <-1 and malnourished at HAZ −2: orange box: 3-year HAZ score −1 to −2; red box: 3-year HAZ score < −2. X-axis indicates number of *Cryptosporidium* infectio**ns**

**Fig S8.**
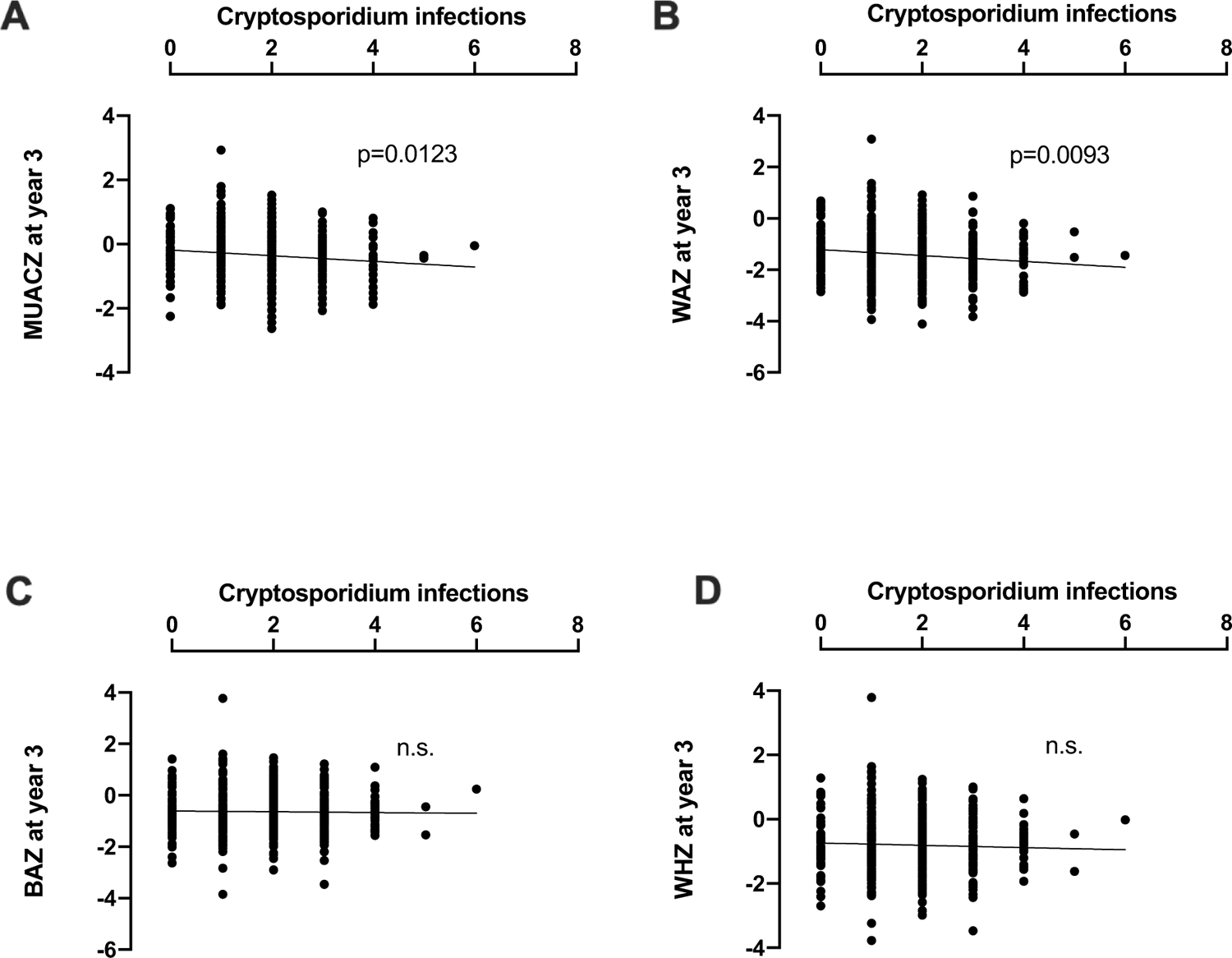
Cryptosporidiosis was associated with chronic but not acute malnutrition at year 3. Graphs show results from a simple linear regression with each symbol representing a single child X-axis indicates number of *Cryptosporidium* infections A) Y-axis MUACZ circumference of the mid-upper arm (muscle wasting) B) Y-axis WHZ score (low weight for height (wasting) a measure of acute malnutrition C) Y axis WAZ score (low weight for age) a measure of acute and chronic malnutrition and D) BAZ (body mass index for age)

**Fig S9.**
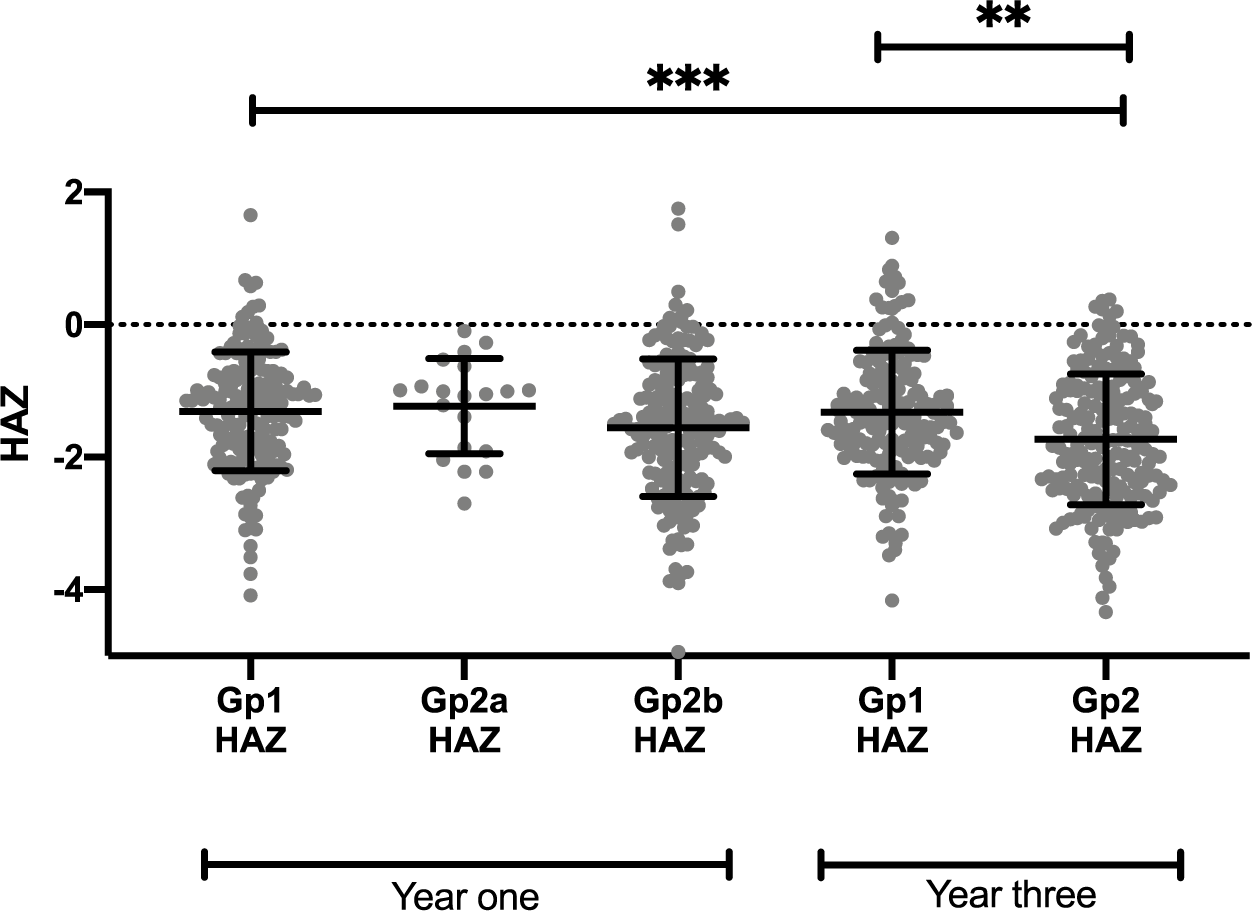
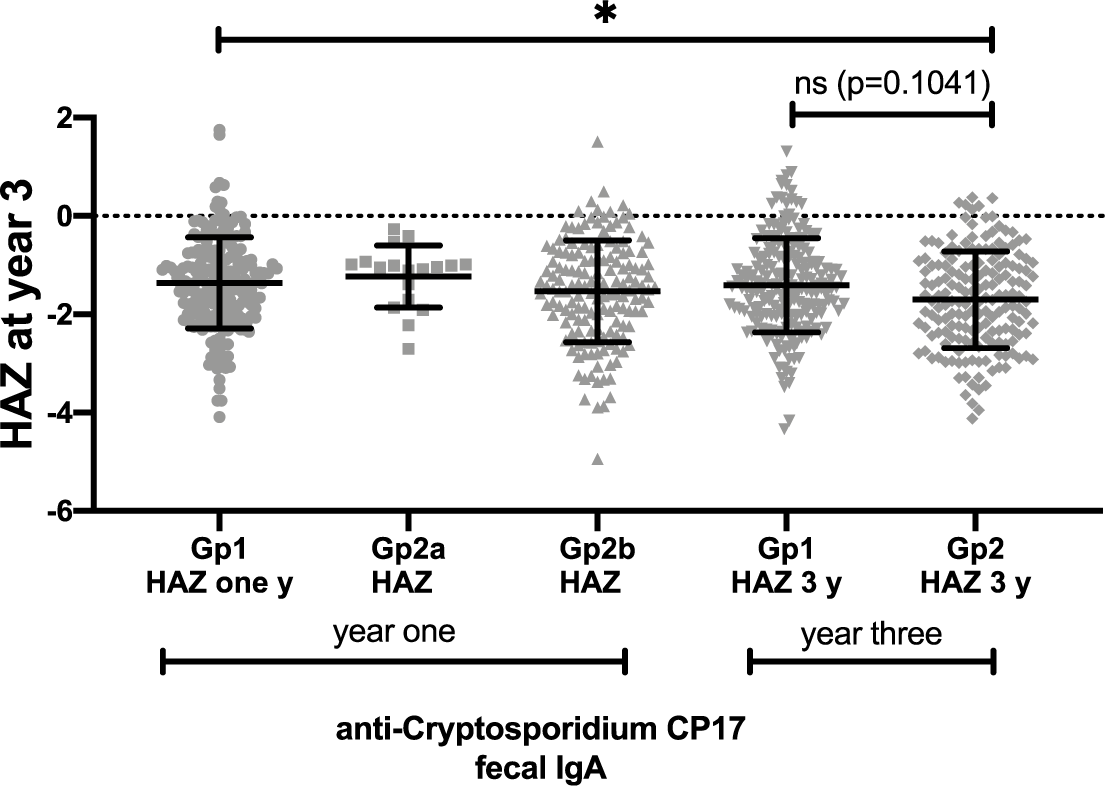
Low anti-Cryptosporidium IgA levels after an infection were associated with a subsequent increase in cryptosporidiosis-associated growth-faltering. Bar graphs (indicating data mean ± standard deviation) with individual data points. Each symbol on the box plot represents a child. On the X-axis values are shown for children in Groups 1 and 2 in both year one and year 3. Groups on the X-axis refers to the values obtained the end of the first and third years of life. Y-axis represents the growth faltering (HAZ). Horizontal bars represent the result of a non-parametric Kruskal-Wallis test *** indicates p<0.001 ** indicates p<0.01 * p<0.05 A) Group 1 children had higher than average levels of fecal anti-IgA Cp23. Group2a children were negative by diagnostic surveillance by qPCR and anti-Cp23 antibodies at year one Group2b were positive by qPCR but had nevertheless low levels of anti-Cp23 antibodies. B) Group 1 children had higher than average levels of fecal anti-IgA Cp17. Group2a children were negative by diagnostic surveillance by qPCR and anti-Cp17 antibodies at year one Group2b were positive by qPCR but had nevertheless low levels of anti-Cp17 antibodies.

**Table S1.** Symptomatic and asymptomatic samples collected during year 3

| Sample Type | Total<br>Diarrhea<br>Recorded | Samples<br>Collected | Samples<br>not<br>Collected | RT_QPCR<br>Assay | Sample not<br>Assayed |
| --- | --- | --- | --- | --- | --- |
| **Year 2** |  |  |  |  |  |
| **Symptomatic** | 3763 | 3148 (83.66%) | **615** | 3132 | **16 (0.5%)** |
| **Sub-clinical\*** | NA | 15312 (97.58%) | **318** | 14638 | **674 (4.4%)** |
| **Year3** |  |  |  |  |  |
| **Symptomatic** | 3578 | 3148 (83.96%) | **574** | 2978 | **26 (0.9%)** |
| **Sub-clinical\*** | NA | 14473 (97.94%) | **305** | 13806 | **667(4.6%)** |
| **\*Monthly Stool Samples** |  |  |  |  |  |

**Table S2.** Multivariable analysis of total all cause diarrhea and HAZ at year 3

| Parameter | Effect (95% Confidence Interval) | P Value |
| --- | --- | --- |
| All cause diarrhea<sup>#</sup> | -0.008 (-0.023, 0.008) | 0.3357 |
| Child LAZ at Birth | 0.245 (0.151, 0.339) | <0.0001 |
| Maternal Weight | 0.017 (0.007, 0.027) | 0.001 |
| Maternal Height | 0.037 (0.019, 0.056) | 0.0001 |
| Maternal Education | 0.233 (0.020, 0.446) | 0.0323 |
| Household income | 0.001 (0.000, 0.002) | 0.079 |
| Treated water | 0.205 (-0.003, 0.413) | 0.0537 |

<sup>#</sup> diarrheal episodes (all-cause) were not significantly associated with HAZ at year 3

## References

1. Sow SO, Muhsen K, Nasrin D, Blackwelder WC, Wu Y, Farag TH, et al. The Burden of Cryptosporidium Diarrheal Disease among Children & 24 Months of Age in Moderate/High Mortality Regions of Sub-Saharan Africa and South Asia, Utilizing Data from the Global Enteric Multicenter Study (GEMS). Kosek M, editor. PLoS Negl Trop Dis. 2016;10: e0004729. doi:10.1371/journal.pntd.0004729

2. Wang H, Abajobir AA, Abate KH, Abbafati C, Abbas KM, Abd-Allah F, et al. Global, regional, and national under-5 mortality, adult mortality, age-specific mortality, and life expectancy, 1970–2016: a systematic analysis for the Global Burden of Disease Study 2016. Lancet. 2017;390: 1084–1150. doi:10.1016/S0140-6736(17)31833-0

3. Korpe PS, Valencia C, Haque R, Mahfuz M, McGrath M, Houpt E, et al. Epidemiology and Risk Factors for Cryptosporidiosis in Children from Eight Low-income Sites: Results from the MAL-ED Study. Clin Infect Dis. 2018; doi:10.1093/cid/ciy355

4. Haque R, Mondal D, Karim A, Molla IH, Rahim A, Faruque ASG, et al. Prospective case-control study of the association between common enteric protozoal parasites and diarrhea in Bangladesh. Clin Infect Dis. 2009;48: 1191–1197. doi:10.1086/597580

5. Cama Va, Ross JM, Crawford S, Kawai V, Chavez-Valdez R, Vargas D, et al. Differences in clinical manifestations among Cryptosporidium species and subtypes in HIV-infected persons. J Infect Dis. 2007;196: 684–91. doi:10.1086/519842

6. Chalmers RM, Davies AP, Tyler K. Cryptosporidium. Microbiol (United Kingdom). 2019;165: 500–502. doi:10.1099/mic.0.000764

7. Chavez MA, White AC. Novel treatment strategies and drugs in development for cryptosporidiosis. Expert Review of Anti-Infective Therapy. Taylor and Francis Ltd; 2018. pp. 655–661. doi:10.1080/14787210.2018.1500457

8. Korpe P, Haque R, Gilchrist CA, Valencia C, Niu F, Lu MM, et al. Natural History of Cryptosporidiosis in a Longitudinal Study of Slum-Dwelling Bangladeshi Children: Association with Severe Malnutrition. PLoS Negl Trop Dis. 2016;10: e0004564. doi:10.1371/journal.pntd.0004564

9. Kattula D, Jeyavelu N, Prabhakaran AD, Premkumar PS, Velusamy V, Venugopal S, et al. Natural History of Cryptosporidiosis in a Birth Cohort in Southern India. Clin Infect Dis. 2017;64: 347–354. doi:10.1093/cid/ciw730

10. Khalil IA, Troeger C, Rao PC, Blacker BF, Brown A, Brewer TG, et al. Morbidity, mortality, and long-term consequences associated with diarrhoea from Cryptosporidium infection in children younger than 5 years: a meta-analyses study. Lancet Glob Heal. 2018;6: e758– e768. doi:10.1016/S2214-109X(18)30283-3

11. Krumkamp R, Aldrich C, Maiga-Ascofare O, J M, Rakotozandrindrainy N, Borrmann S, et al. Transmission of Cryptosporidium spp. among human and animal local contact networks in sub-Saharan Africa: a multi-country study. Clin Infect Dis. 2020; ciaa223. Available: https://pubmed.ncbi.nlm.nih.gov/32150243/

12. Eibach D, Krumkamp R, Al-Emran HM, Sarpong N, Hagen RM, Adu-Sarkodie Y, et al. Molecular characterization of Cryptosporidium spp. among children in rural Ghana. PLoS Negl Trop Dis. 2015;9: e0003551. doi:10.1371/journal.pntd.0003551

13. Steiner KL, Ahmed S, Gilchrist CA, Burkey C, Cook H, Ma JZ, et al. Species of Cryptosporidia Causing Subclinical Infection Associated With Growth Faltering in Rural and Urban Bangladesh: A Birth Cohort Study. Clin Infect Dis. 2018;67: 1347–1355. doi:10.1093/cid/ciy310

14. Steiner KL, Kabir M, Hossain B, Gilchrist CA, Ma JZ, Ahmed T, et al. Delayed Time to Cryptosporidiosis in Bangladeshi Children is Associated with Greater Fecal IgA against Two Sporozoite-Expressed Antigens. Am J Trop Med Hyg. 2021;104: 229–232. doi:10.4269/ajtmh.20-0657

15. Taniuchi M, Sobuz SU, Begum S, Platts-Mills JA, Liu J, Yang Z, et al. Etiology of Diarrhea in Bangladeshi Infants in the First Year of Life Using Molecular Methods. J Infect Dis. 2013;e-pub ahea: jit507-. doi:10.1093/infdis/jit507

16. Lemieux MW, Sonzogni-Desautels K, Ndao M. Lessons learned from protective immune responses to optimize vaccines against cryptosporidiosis [Internet]. Pathogens. MDPI AG; 2018. doi:10.3390/pathogens7010002

17. Sateriale A, Šlapeta J, Baptista R, Engiles JB, Gullicksrud JA, Herbert GT, et al. A Genetically Tractable, Natural Mouse Model of Cryptosporidiosis Offers Insights into Host Protective Immunity. Cell Host Microbe. 2019;26: 135–146.e5. doi:10.1016/j.chom.2019.05.006

18. Checkley W, White AC, Jaganath D, Arrowood MJ, Chalmers RM, Chen X-MM, et al. A review of the global burden, novel diagnostics, therapeutics, and vaccine targets for cryptosporidium. Lancet Infect Dis. 2015;15: 85–94. doi:10.1016/S1473-3099(14)70772-8

19. Tzipori S, Roberton D, Chapman C. Remission of diarrhoea due to cryptosporidiosis in an immunodeficient child treated with hyperimmune bovine colostrum. Br Med J. 1986;293: 1276–7. Available: http://www.pubmedcentral.nih.gov/articlerender.fcgi?artid=1342109&tool=pmcentrez&rendertype=abstract

20. Steiner KL, Kabir M, Priest JW, Hossain B, Gilchrist CA, Cook H, et al. Fecal IgA against a sporozoite antigen at 12 months is associated with delayed time to subsequent cryptosporidiosis in urban Bangladesh: a prospective cohort study. Clin Infect Dis. 2019; doi:10.1093/cid/ciz430

21. Steiner KL, Kabir M, Priest JW, Hossain B, Gilchrist CA, Cook H, et al. Fecal immunoglobulin a against a sporozoite antigen at 12 months is associated with delayed time to subsequent cryptosporidiosis in urban Bangladesh: A prospective cohort study. Clin Infect Dis. 2020;70: 323–326. doi:10.1093/cid/ciz430

22. Ajjampur SSR, Sarkar R, Sankaran P, Kannan A, Menon VK, Muliyil J, et al. Symptomatic and asymptomatic *Cryptosporidium* infections in children in a semi-urban slum community in southern India. Am J Trop Med Hyg. 2010;83: 1110–5. doi:10.4269/ajtmh.2010.09-0644

23. Okhuysen PC, Chappell CL, Sterling CR, Jakubowski W, DuPont HL. Susceptibility and serologic response of healthy adults to reinfection with Cryptosporidium parvum. Infect Immun. 1998;66: 441–443. doi:10.1128/iai.66.2.441-443.1998

24. Guerrant DI, Moore SR, Lima AAM, Patrick PD, Schorling JB, Guerrant RL. Association of early childhood diarrhea and cryptosporidiosis with impaired physical fitness and cognitive function four-seven years later in apoor urban community in northeast Brazil. Am J Trop Med Hyg. 1999;61: 707–713. doi:10.4269/ajtmh.1999.61.707

25. Checkley W, Gilman RH, Epstein LD, Suarez M, Diaz JF, Cabrera L, et al. Asymptomatic and symptomatic cryptosporidiosis: their acute effect on weight gain in Peruvian children. Am J Epidemiol. 1997;145: 156–63. doi:10.1093/oxfordjournals.aje.a009086

26. Schnee AE, Haque R, Taniuchi M, Uddin J, Alam M, Liu J, et al. Identification of etiology-specific diarrhea associated with linear growth faltering in Bangladeshi infants. Am J Epidemiol. 2018;187: 2210–2218. doi:10.1093/aje/kwy106

27. de Onis M, Branca F. Childhood stunting: A global perspective. Matern Child Nutr. 2016;12: 12–26. doi:10.1111/mcn.12231

28. Guide I. Interpretation Guide. Nutr Landacape Inf Syst. 2012; 1–51. doi:10.1159/000362780.Interpretation

29. Liu J, Platts-Mills JA, Juma J, Kabir F, Nkeze J, Okoi C, et al. Use of quantitative molecular diagnostic methods to identify causes of diarrhoea in children: a reanalysis of the GEMS case-control study. Lancet. 2016;388: 1291–1301. doi:10.1016/S0140-6736(16)31529-X

30. Taniuchi M, Sobuz SU, Begum S, Platts-Mills JA, Liu J, Yang Z, et al. Etiology of diarrhea in Bangladeshi infants in the first year of life analyzed using molecular methods. J Infect Dis. 2013;208: 1794–802. doi:10.1093/infdis/jit507

31. Platts-Mills JA, Liu J RE, et al. Aetiology, burden and clinical characteristics of diarrhea in children in low-resource settings using quantitative molecular diagnostics: results from the MAL-ED cohort study. Lancet Glob Heal. 2018;in press.

32. Cama VA, Bern C, Roberts J, Cabrera L, Sterling CR, Ortega Y, et al. Cryptosporidium species and subtypes and clinical manifestations in children, Peru. Emerg Infect Dis. 2008;14: 1567–74. doi:10.3201/eid1410.071273

33. Chappell CL, Okhuysen PC, Langer-Curry RC, Akiyoshi DE, Widmer G, Tzipori S. Cryptosporidium meleagridis: Infectivity in healthy adult volunteers. Am J Trop Med Hyg. 2011;85: 238–242. doi:10.4269/ajtmh.2011.10-0664

34. Ajjampur SSR, Liakath FB, Kannan A, Rajendran P, Sarkar R, Moses PD, et al. Multisite study of cryptosporidiosis in children with diarrhea in India. J Clin Microbiol. 2010;48: 2075–81. doi:10.1128/JCM.02509-09

35. Dong S, Yang Y, Wang Y, Yang D, Yang Y, Shi Y, et al. Prevalence of Cryptosporidium Infection in the Global Population: A Systematic Review and Meta-analysis. Acta Parasitol. 2020; doi:10.2478/s11686-020-00230-1

36. Hossain MJ, Saha D, Antonio M, Nasrin D, Blackwelder WC, Ikumapayi UN, et al. Cryptosporidium infection in rural Gambian children: Epidemiology and risk factors. Bartelt LA, editor. PLoS Negl Trop Dis. 2019;13: e0007607. doi:10.1371/journal.pntd.0007607

37. Bushen OY, Kohli A, Pinkerton RC, Dupnik K, Newman RD, Sears CL, et al. Heavy cryptosporidial infections in children in northeast Brazil: comparison of Cryptosporidium hominis and Cryptosporidium parvum. Trans R Soc Trop Med Hyg. 2007;101: 378–84. doi:10.1016/j.trstmh.2006.06.005

